# Primulagenin A is a potent inverse agonist of the nuclear receptor RAR-related orphan receptor gamma (RORγ)

**DOI:** 10.1101/2025.04.01.646598

**Authors:** Patrik F. Schwarz, Alexander F. Perhal, Teresa Preglej, Lina Breit, Klaus G. Schmetterer, Ulrike Grienke, Jasmin Janneschütz, Ya Chen, Natacha Rochel, Johannes Kirchmair, Nina Schützenmeister, Judith M. Rollinger, Michael Bonelli, Verena M. Dirsch

## Abstract

Throughout history, herbal medicines and natural products have played a crucial role as therapeutics for humans, yet their molecular mechanisms of action often remain elusive. Here, we ask whether primulagenin A (PGA) from the traditionally used herbal substance Primula root, acts *via* the nuclear receptor RORγ, a key regulator of pro-inflammatory Th17 cells, which are linked to autoimmune diseases like psoriasis. Luciferase assays revealed a high potency (IC_50_ ~100 nM) and efficacy (I_max_ ~ 90%) of PGA as an inverse agonist of RORγ. To ensure sufficient supply, we established methods to isolate and synthesize PGA. Its binding to the human RORγ ligand binding domain was confirmed by nano differential scanning fluorimetry, and a structure-activity relationship was proposed by docking and site-directed mutagenesis. PGA downregulated RORγ target gene expression and inhibited murine and human Th17 differentiation in a concentration-dependent manner. It also reduced the proportion of IL-17A-producing Th17 cells. In this work, we identify PGA as a new, potent, and efficacious inverse agonist of RORγ, with potential for modulating immune responses in inflammatory and autoimmune diseases.

## Introduction

For centuries, humans have relied on medicinal plants as a fundamental source of therapeutic compounds[1]. The importance of “unlocking nature’s pharmacy” is underpinned by the number of newly approved drugs that are either natural products, derived from natural products or “defined mixtures” that sum up to 23.5% of all drugs approved between 01/1981 and 09/2019[2]. Prominent examples of FDA-approved natural products include digoxin[3], morphine[4], and colchicine[5]. In contrast to these well-characterized molecules, the precise molecular mechanism of many other purified natural products is hard to pin down, even if cellular (e.g., antiproliferative) effects appear promising[6]. When it comes to herbal medicinal products or herbal substances, there is often a strong contrast between their successful use for centuries and the elusiveness of their pharmacological mode of action[7, 8]. For instance, the rhizomes and roots of cowslip (*Primula veris* L.) or oxlip (*Primula elatior* (L.) Hill) – known as “Primula root” or “*Primulae radix*”– are traditionally used to treat cough and cold. In Europe, several herbal preparations containing Primulae radix in liquid and solid dosage forms are marketed as herbal medicinal products against respiratory infections. Still, studies providing a solid rationale for this use are scarce[9]. In 2005, Nauert *et al.* reported that a fluid extract from Primulae radix concentration-dependently reduced the lipopolysaccharide-induced interleukin (IL)-8 release from primary monocytes. In contrast, a fluid extract from thyme hardly affected the release of this cytokine (reviewed in [10]). Interestingly, clinical trials examining the effects of fixed combinations of both herbal substances found favorable effects for treating acute bronchitis[11–13]. Despite these promising findings, a molecular target remained unknown.

The RAR-related orphan receptors (RORs) are a subfamily of nuclear receptors (NRs) consisting of RORα[14], RORβ[15], RORγ[16], and various isoforms of these proteins[17]. In 2006, RORγt, the RORγ isoform exclusively expressed in cells of the immune system, was identified as the crucial transcription factor driving the differentiation of naïve CD4^+^ T cells into pro-inflammatory T helper (Th)-17 cells[18]. Th17 cells possess a dichotomous nature. They secrete cytokines like IL-17A, IL-17F, and IL-22 and act as host defenders against bacterial and fungal infections. Furthermore, they play a significant role in autoimmune diseases like psoriasis, rheumatoid arthritis (RA), inflammatory bowel disease (IBD), and multiple sclerosis (MS)[19]. As RORγt is an important transcription factor for Th17 cell differentiation and function of differentiated Th17 cells[20, 21], inhibition of this NR was brought up as a potential therapy option for these autoimmune diseases[22]. Depending on whether RORγ is considered unliganded or liganded inside cells, RORγ inhibitors can be classified as inverse agonists or antagonists, respectively[23]. Given that apo-RORγ has demonstrated the capacity to adopt an active conformation in the absence of any ligands, RORγ inhibitors will be referred to as inverse agonists in this study[24]. Examples of such inverse agonists range from synthetic compounds like SR2211[25] to a myriad of natural products, including triterpenoids like ursolic acid (UA) and oleanolic acid (OA)[26]. Interestingly, *Primula elatior*, specifically, contains oleanane-type triterpene saponins that are hydrolysed to primulagenin A (PGA) under acidic conditions[9, 27, 28]. Here, we present PGA as a new and highly active inverse agonist of RORγ, highlighting its promise for therapeutic immunomodulation through inhibition of the pro-inflammatory Th17 response.

## Results

### PGA is a new oleanane-type triterpenoid RORγ inverse agonist with high potency and efficacy

Due to its structural similarity to the known inverse agonist OA, PGA (**Figure 1A**) was probed for its ability to inhibit RORγ in a Gal4 luciferase assay. **Figure 1B** depicts the results of the initial screening. PGA led to a concentration-dependent decrease in transcriptional activity of RORγ-Gal4 in HEK293 cells and yielded an IC_50_ value of approx. 75 nM and an I_max_ of 0.319-fold (68% inhibition) at 10 µM. These results were confirmed using a full-length RORγ luciferase assay (**Figure 1C**), in which PGA showed an IC_50_ value of 119 nM and an I_max_ value of 0.120-fold (88% inhibition) at 10 µM.

**Figure 1:**
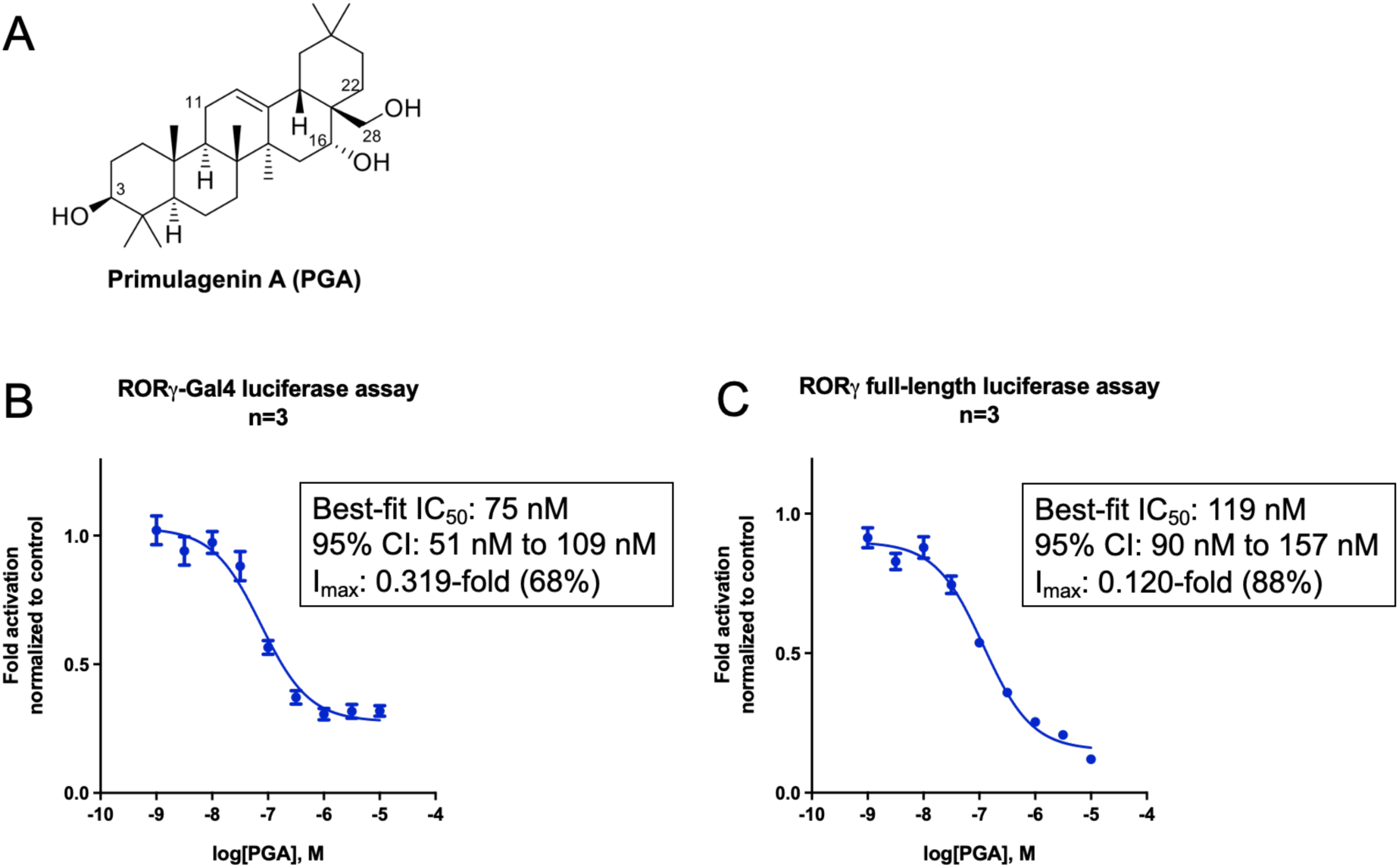
PGA acts as a highly potent and efficacious RORγ inverse agonist. **A** Chemical structure of PGA. **B** Concentration-response curve of PGA showing a concentration-dependent inhibition of the transcriptional activity of RORγ in a RORγ-Gal4 luciferase assay. HEK293 cells were co-transfected with RORγ-Gal4, an UAS luciferase reporter, and eGFP. The cells were then treated with PGA at the indicated concentration. The RLU/RFU ratio was measured after an incubation time of 18 hours and normalized to the vehicle control. Mean ± SEM, n=3 in technical quadruplicates. **C** Concentration-response curve of PGA showing a concentration-dependent inhibition of the transcriptional activity of RORγ in a full-length RORγ luciferase assay. HEK293 cells were co-transfected with full-length RORγ, a luciferase reporter under the control of RORE, and eGFP, and treated with PGA at the indicated concentration. The RLU/RFU ratio was measured after an incubation time of 18 hours and normalized to the vehicle control. Mean ± SEM, n=3 in technical quadruplicates.

To exclude the possibility that cytotoxicity interfered with the results, a resazurin conversion assays in HEK293 cells was performed that revealed no cytotoxic effects (**Figure S 2A**). Finally, additional NR-luciferase assays were conducted to confirm the selectivity of PGA for RORγ over other NRs. The following NRs were chosen for selectivity screenings as they are targeted by published triterpenoid RORγ inverse agonists like UA and OA: farnesoid X receptor (FXR), liver X receptor (LXR)α/β, peroxisome proliferator–activated receptor (PPAR)β/γ, murine retinoic acid receptor (mRAR)α, and retinoid X receptor (RXR)α/β (reviewed in[29, 30]). To assess the potential selectivity of PGA within the ROR subfamily, it was tested on RORα and RORβ as well. Off-target effects of PGA were not detected on any of the receptors tested (**Figure S 3**). To summarize, PGA was identified as a new triterpenoid RORγ inverse agonist with high potency and efficacy in cellular *in vitro* luciferase assays. Cytotoxicity was not detected, and PGA showed selectivity for RORγ over other NRs usually co-modulated by triterpenoids and over RORα and RORβ.

### PGA is accessible in large quantities and high purity via isolation from Primulae radix or by organic synthesis

For further pharmacological profiling, PGA was isolated from a suitable natural source, as it was not commercially available. Therefore, the roots of *Primula* sp. were selected for large scale phytochemical processing and subsequent isolation of PGA. A simplified overview of this process is depicted in **Figure 2A**. A hydrolyzed and sapogenin-enriched extract of Primulae radix (PRA_A) was fractionated *via* flash chromatography and size exclusion chromatography to obtain PRA_A 2.4, which was further subjected to semi-preparative supercritical fluid chromatography (SFC-15) to obtain seven subfractions (PRA_A 3.1-3.7). The bio-guided phytochemical workup clearly pointed towards PRA_A 3.5 as the major RORγ inhibiting component (**Figure 2B**), which revealed to be PGA with a purity of > 95% (**Table S 1**). Interestingly, it was found that the methanolic Primulae radix extract, from which PGA was isolated after hydrolysis, and a commercially available cough syrup share a highly similar saponin profile (data not shown).

**Figure 2:**
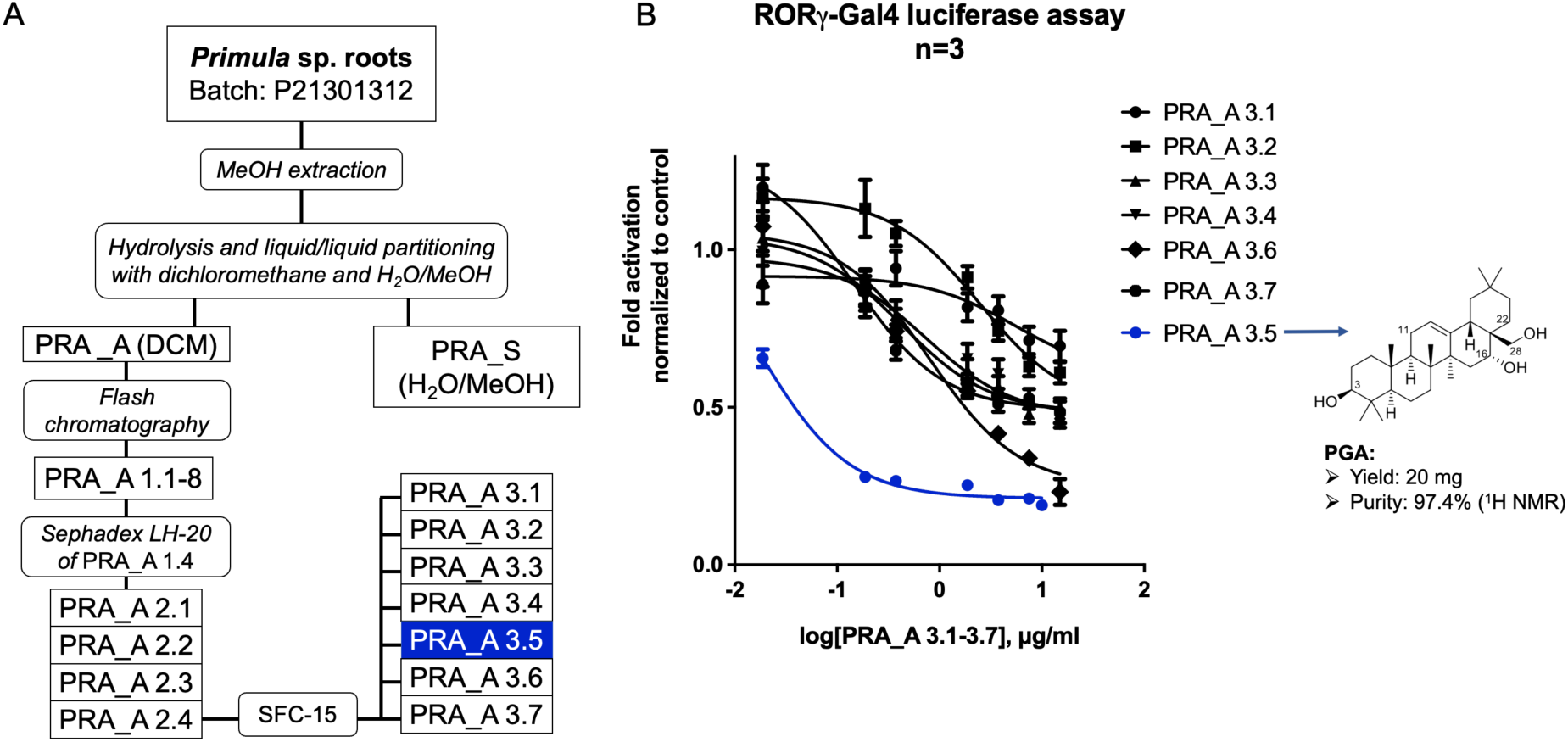
PGA was isolated from Primulae radix in high quantity and purity. **A** Simplified overview of the isolation process. **B** Concentration-response curve of the subfractions PRA_A 3.1-3.7 reveal that PRA_A 3.5 (later identified as PGA) has the highest inhibitory activity on RORγ in a RORγ-Gal4 luciferase assay. HEK293 cells were co-transfected with RORγ-Gal4, an UAS luciferase reporter, and eGFP, and treated with the extracts at the indicated concentration. The RLU/RFU ratio was measured after an incubation time of 18 hours and normalized to the vehicle control. Mean ± SEM, n=3 in technical quadruplicates.

To attain an alternate pathway towards PGA, a synthetic route starting from commercially available echinocystic acid was established (described in **“Synthesis of primulagenin A”**; ^1^H- and ^13^C-NMR spectra are shown in **Figure S 4**). The identity and purity of both, the isolated and synthesized PGA were confirmed (**Table S 1**) and thus used for all subsequent experiments. In conclusion, two independent routes to replenish PGA for further biological studies were successfully established.

### Structure-activity relationship of PGA and related oleanane-type triterpenoids reveal the importance of C-16 and C-28 for the activity of these compounds

Compared to the potency and efficacy of PGA, the reported inverse agonistic activity of the triterpenoid OA on RORγ (IC_50_ = 8.589 µM in a luciferase assay[31]) seemed underwhelming. Hence, the luciferase assay for OA was repeated and confirmed previous findings (**Table 1**). Both, PGA, and OA are oleanane-type triterpenoids that share a high structural similarity. They only differ with respect to the oxidation state on C-atom 28 (−CH_2_OH (PGA) vs. −CO_2_H (OA); **Table 1** – “R1”) and C-atom 16 (−OH (PGA) vs. −H (OA); **Table 1** – “R2”). A lower oxidation state of R1 and the presence of an oxygen atom on R2 appeared to be the critical factors for the superior activity of PGA over OA. To confirm this hypothesis, structurally related triterpenoids from our in-house pure compound database were tested that differed in these groups in RORγ-Gal4 and RORγ full-length luciferase assays. Primulagenin D (PGD; R1 = −CHO, R2 = −OH), echinocystic acid (EA; R1 = −CO_2_H, R2 = −OH), and β-amyrin (β-Amy; R1 = −CH_3_, R2 = −H) were selected to complement the data. Indeed, an increase in the oxidation state of R1 from an alcohol (PGA) to an aldehyde (PGD) to a carboxylic acid moiety (EA) led to a consistent decrease in potency and efficacy (**Table 1** and **Figure 3A, B**).

**Table 1:** SAR of primulagenin A (PGA), primulagenin D (PGD), echinocystic acid (EA), erythrodiol (ED), oleanolic aldehyde (OAL), oleanolic acid (OA), and β-amyrin (β-Amy). IC50 and Imax values of these triterpenoids are depicted. Data were collected using RORγ-Gal4 and full-length RORγ luciferase assays. n=3 in all cases.

| Tri-<br>terpenoid | R<sup>1</sup> | R<sup>2</sup> | Best-fit IC<sub>50</sub><br>(μM) | IC<sub>50</sub> 95% CI<br>(μM) | I<sub>max</sub> at 10 μM |
| --- | --- | --- | --- | --- | --- |
| **RORγ-Gal4 luciferase assay** |  |  |  |  |  |
| **PGA** | -CH<sub>2</sub>OH | -OH | 0.099 | 0.068 – 0.143 | 0.228-fold (77% inh.) |
| **PGD** | -CHO | -OH | 0.542 | 0.335 – 0.889 | 0.290-fold (71% inh.) |
| **EA** | -CO<sub>2</sub>H | -OH | 2.127 | 1.082 – 4.474 | 0.349-fold (65% inh.) |
| **ED** | -CH<sub>2</sub>OH | -H | 0.456 | 0.214 – 0.968 | 0.627-fold (37% inh.) |
| **OAL** | -CHO | -H | 1.739 | 0.381 – 21.300 | 0.653-fold (35% inh.) |
| **OA** | -CO<sub>2</sub>H | -H | 3.625 | 0.922 – 27.260 | 0.429-fold (57% inh.) |
| **β-Amy** | -CH<sub>3</sub> | -H | 6.666 | 0.751 – n.d. | 0.674-fold (33% inh.) |
| **ROR<sub>γ</sub> full-length luciferase assay** |  |  |  |  |  |
| **PGA** | -CH<sub>2</sub>OH | -OH | 0.165 | 0.139 – 0.197 | 0.086-fold (91% inh.) |
| **PGD** | -CHO | -OH | 0.717 | 0.463 – 1.113 | 0.246-fold (75% inh.) |
| **EA** | -CO<sub>2</sub>H | -OH | 1.556 | 1.010 – 2.436 | 0.313-fold (69% inh.) |
| **ED** | -CH<sub>2</sub>OH | -H | 0.143 | 0.037 – 0.622 | 0.778-fold (22% inh.) |
| **OAL** | -CHO | -H | Ambiguous | Ambiguous | 0.964-fold (4% inh.) |
| **OA** | -CO<sub>2</sub>H | -H | 2.988 | 1.033 – 10.230 | 0.464-fold (54% inh.) |
| **β-Amy** | -CH<sub>3</sub> | -H | 14.770 | 1.128 – n.d. | 0.775-fold (23% inh.) |

**Figure 3:**
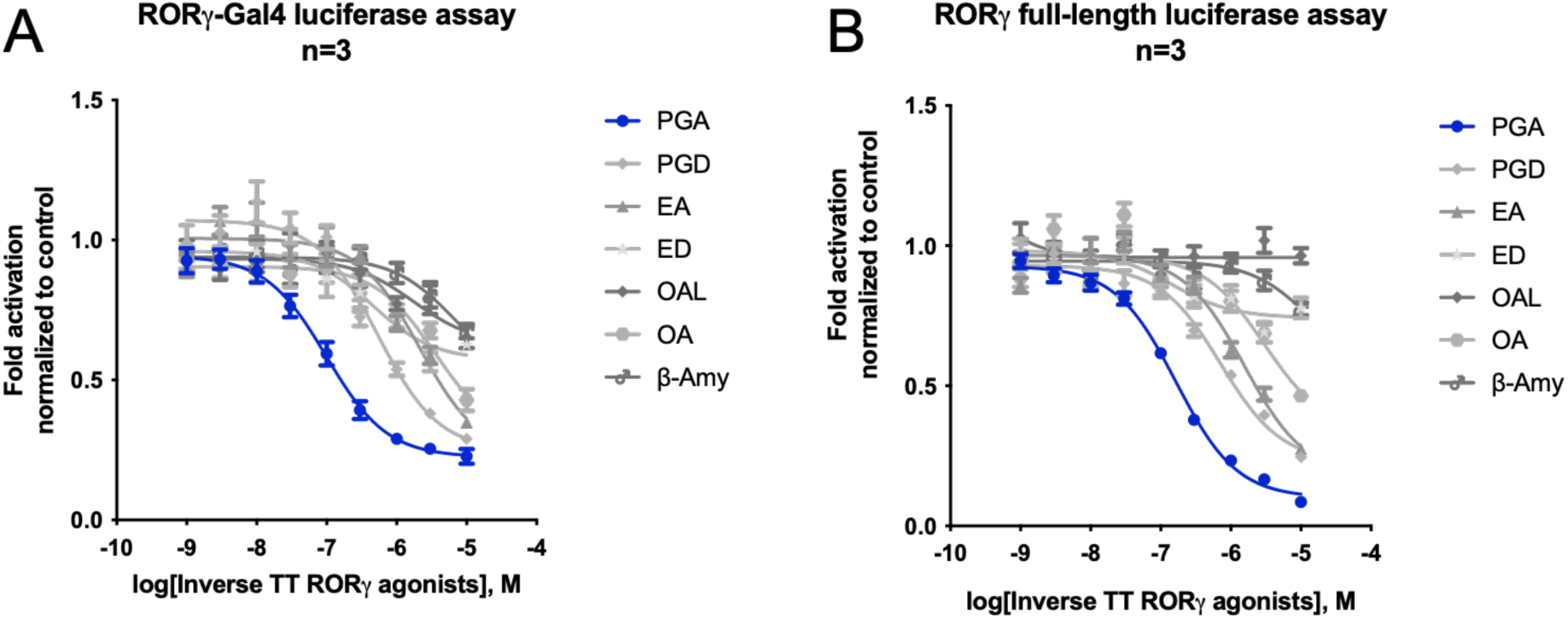
Differing activities of PGA and structurally related triterpenoids as RORγ inverse agonists. **A** Concentration-response curves of PGA and structurally related triterpenoids using a RORγ-Gal4 luciferase assay. HEK293 cells were co-transfected with RORγ-Gal4, an UAS luciferase reporter, and eGFP, and treated with the triterpenoids at the indicated concentration. The RLU/RFU ratio was measured after an incubation time of 18 hours and normalized to the vehicle control. Mean ± SEM, n=3 in technical quadruplicates. **B** Concentration-response curves of PGA and structurally related triterpenoids using a full-length RORγ luciferase assay. HEK293 cells were co-transfected with full-length RORγ, a luciferase reporter under the control of RORE, and eGFP, and treated with the triterpenoids at the indicated concentration. The RLU/RFU ratio was measured after an incubation time of 18 hours and normalized to the vehicle control. Mean ± SEM, n=3 in technical quadruplicates.

The lack of an −OH group on R2 further decreased both, potency and efficacy (e.g., EA vs. OA, **Table 1** and **Figure 3A, B**). Lastly, a CH_3_-group on R1 and a H-atom on R2 (β-Amy) abolished RORγ activity (**Table 1** and **Figure 3A, B**). Interestingly, erythrodiol (ED; R1 = CH_2_OH, R2 = −H; synthesis described in **“Synthesis of erythrodiol”** and ^1^H- and ^13^C-NMR spectra shown in **Figure S 5**), showed a favorable potency but a loss of efficacy (**Table 1** and **Figure 3A, B**), underpinning the importance of an −OH group on R2. This was even more pronounced in case of the newly synthesized compound oleanolic aldehyde (OAL; R1 = −CHO, R2 = −H; synthesis described in **“Synthesis of oleanolic aldehyde”** and ^1^H- and ^13^C-NMR spectra shown in **Figure S 6**), which was completely inactive on full-length RORγ (**Table 1** and **Figure 3A, B**). To test whether the superiority of PGA was statistically significant, we compared the best-fit log IC_50_ values of PGA with all other structurally related triterpenoids listed in **Table 1** using an extra sum-of-squares F test. Indeed, PGA was statistically significantly different from the other triterpenoids in terms of its log IC_50_ value (**Table S 2**). Importantly, with the same test, the difference of the log IC_50_ value of PGA from the highly active ursane-type triterpenoid RORγ inverse agonist UA was confirmed to be statistically significant[32] (**Figure S 7**). No cytotoxic effects of any of the compounds at 10 µM (**Figure S 2B**) were observed and we confirmed a high purity for all of them (see **Table S 1** for isolated or synthesized triterpenoids and **Table S 7** for commercially obtained ones). All chemical structures discussed in this work are summarized in **Figure S 1**. Taken together, a low oxidation state on R1 combined with the presence of an −OH group on R2 led to the highest inverse agonistic activity of oleanane-type triterpenoids on RORγ. Moreover, it was shown for the first time that the oleanane-type triterpenoids PGD, EA, and ED are also inverse agonists of RORγ.

### Molecular docking and site-directed mutagenesis suggest unique and distinct protein-ligand interaction patterns for PGA and OA

First, to verify direct binding of PGA to the human RORγ (hRORγ) LBD, nano differential scanning fluorimetry (nanoDSF) was employed[33]. As shown in **Figure 4A**, PGA stabilized the hRORγ LBD to a greater extent (melting temperature of approx. 53.0 °C) compared to the inverse RORγ agonist T0901317 (approx. 50.7 °C)[34]. Next, to understand the structural basis for the differing levels of potency and efficacy between PGA and the published oleanane-type triterpenoid OA on RORγ, docking and site-directed mutagenesis experiments were performed. **Figure 4B** shows the docking poses derived for PGA and OA bound to the RORγ LBD. The 28-OH group of PGA was predicted to form a direct hydrogen bond to glutamine 286 (Q286) of the RORγ LBD, while OA was unlikely to form such an interaction. Consistent with this prediction, the Q286T mutation led to a significant reduction of the activity of PGA compared to its activity on the wild-type (WT) NR while the activity of OA was only slightly (but still significantly) reduced (**Figure 4C**). Notably, PGA was predicted to form an additional hydrogen bond with the backbone of F377 using its 16-OH group, a group that OA lacks.

**Figure 4:**
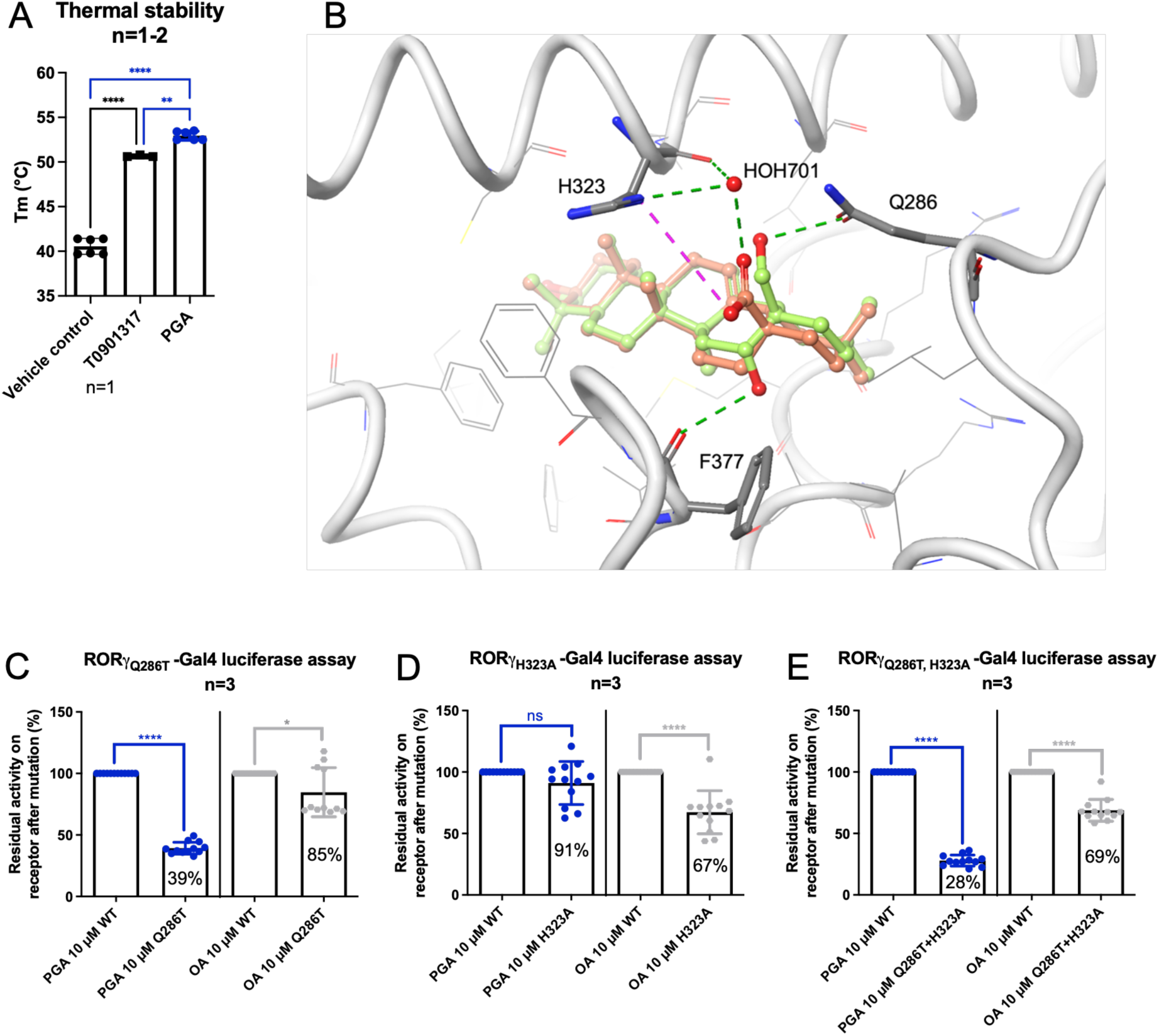
PGA directly binds to the hRORγ LBD, and both PGA and OA are predicted to form partially distinct protein-ligand interactions with the RORγ LBD. **A** Thermal stabilization upon binding of T0901317 and PGA to the hRORγ LBD. nanoDSF was used to compare the thermal stability of the purified apo hRORγ LBD and the ligand-bound protein. Mean ± SD, n=1-2 in technical triplicates. One-way ANOVA with Tukey’s test was performed for statistical analysis. **** P ≤ 0.0001, *** P ≤ 0.001. **B** Docking pose illustrating how PGA (green carbon atoms) and OA (orange carbon atoms) are expected to bind to the human RORγ LBD (PDB structure: 6J3N, in gray carbon atoms and tube representation) by molecular docking. Residues predicted to engage in important interactions with the ligands are marked and highlighted by thick tube representations. Predicted hydrogen bonds are indicated by green dashed lines, and the salt bridge by magenta dashed lines. C-E RORγmutant-Gal4 luciferase assays showing the differences in activity between PGA or OA on the WT NR (100% activity) vs. the respective mutant NR (% residual activity). HEK293 cells were co-transfected with the RORγ-Gal4 WT or a mutant, an UAS luciferase reporter, and eGFP, and treated with PGA or OA at 10 µM. The RLU/RFU ratio was measured after an incubation time of 18 hours and normalized to the vehicle control. Mean ± SD, n=3 in technical quadruplicates. Student’s two-tailed t test was performed for statistical analysis. **** P ≤ 0.0001, * P ≤ 0.05, ns P > 0.05.

Furthermore, OA was predicted to form a direct and indirect hydrogen bond (mediated by water molecule 701) with H323 of the RORγ LBD. In contrast, PGA was predicted not to interact with this amino acid. Consistent with these predictions, the H323A mutation left the activity of PGA unaltered. However, the mutation led to a significant decrease in OA’s activity on the NR compared to WT (**Figure 4D**). Of note, the H323A mutation did not affect the activity of OA as much as the Q286T mutation did in the case of PGA. This suggests that the hydrogen bond formed between OA and H323 might indeed be mediated *via* water molecule 701, as predicted. The Q286T+H323A double mutant resulted in similar reductions in activity (**Figure 4E**), supporting the notion that both mutations independently reduce the activity of PGA and OA, respectively, but not synergistically. The “raw” changes in fold activation after PGA or OA treatment of WT vs. RORγ mutants are depicted in **Figure S 8**. From a molecular point of view, the higher activity of PGA compared to the published RORγ inverse agonist OA is predicted to be due to hydrogen bond “anchoring” (with the backbone of F377) within the RORγ LBD, made possible by the additional 16-OH group. Moreover, the energy penalty related to the desolvation of the carboxylic acid moiety in position C-28 could also be a decisive factor for the lower activity of OA compared to PGA. Both predictions align well with our observations regarding the SAR (see previous section). Overall, direct binding of PGA to the hRORγ LBD was confirmed and the likely differences in protein-ligand interaction patterns for PGA and OA within the RORγ LBD were elucidated.

### PGA downregulates the expression of RORγ target genes in different cell lines

To examine whether PGA affects RORγ target gene expression, qPCR experiments were performed using three different cell lines, HepG2, Jurkat T, and EL-4 cells. HepG2 and Jurkat T cells were transiently transfected with human RORγ prior to experiments. EL-4 cells were stably transfected with murine RORγ (see **“Generation of the stable EL-4-mRORγt cell line”** for details). As expected, PGA led to a significant down-regulation of the RORγ target genes *Il17a/f*, *Il23r* (all of them in EL-4-mRORγt cells), *IL-17A* (Jurkat T cells), and glucose-6-phosphatase (*G6PC*; HepG2 cells) (**Figure 5A-C**) while showing no cytotoxic effect in any of the cell lines used (**Figure S 2C-E**).

**Figure 5:**
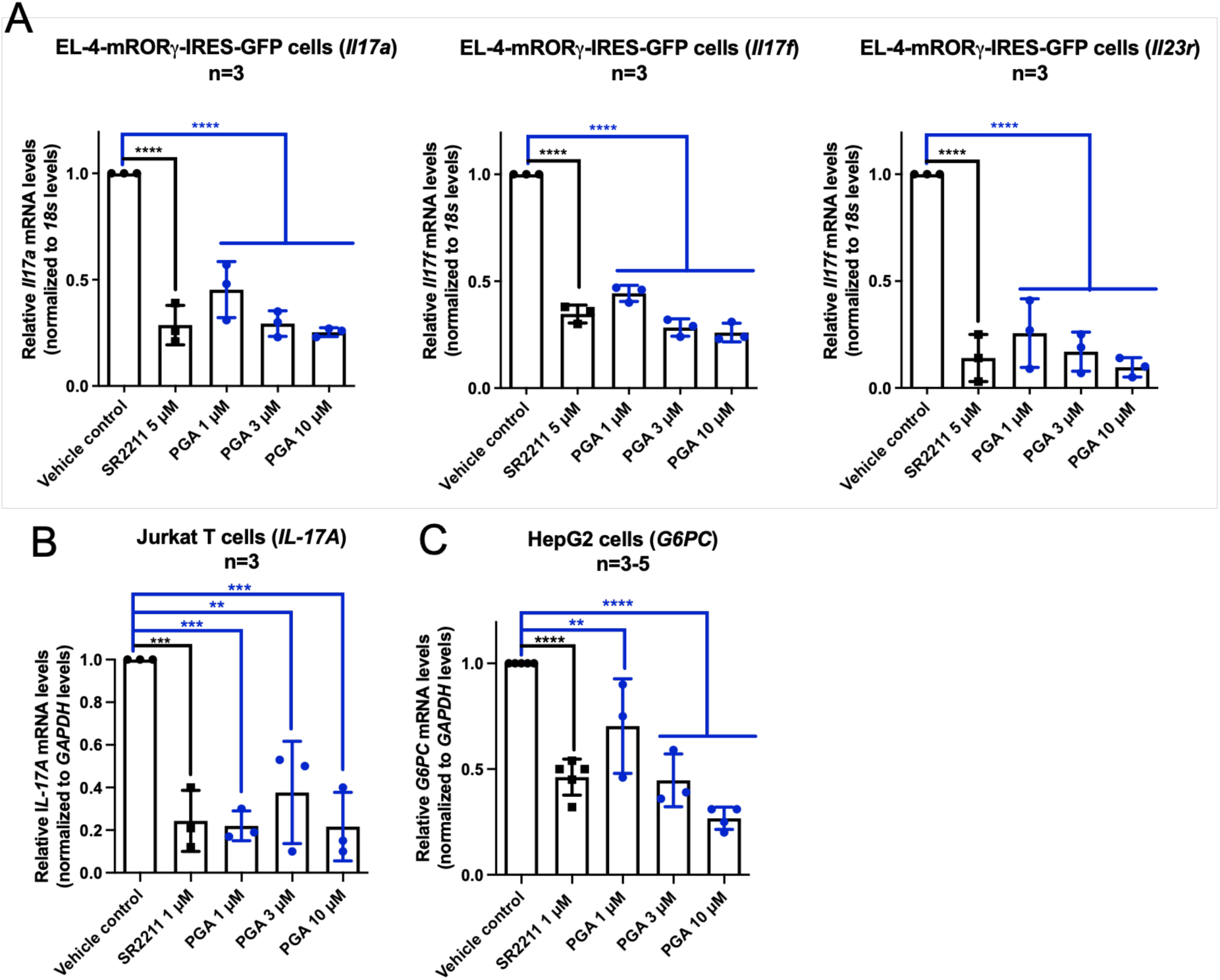
PGA decreases the expression of RORγ target genes in three different cell lines. **A** Decrease of Il17a, Il17f, and Il23r expression in the EL-4-mRORγt cell line upon PGA treatment. 18s was used as a control gene. Mean ± SD, n=3 in technical triplicates. **B** Decrease of IL-17A expression in Jurkat T cells transiently transfected with hRORγ upon PGA treatment. GAPDH was used as a control gene. n=3 in technical triplicates. **C** Decrease of G6PC expression in HepG2 cells transiently transfected with hRORγ upon PGA treatment. GAPDH was used as a control gene. Mean ± SD, n=5 (vehicle control, SR2211), n=4 (PGA 10 µM), n=3 (rest) in technical triplicates. One-way ANOVA with Dunnett’s test was performed for statistical analysis. **** P ≤ 0.0001, *** P ≤ 0.001, ** P ≤ 0.01.

In brief, PGA led to a decreased expression of RORγ target genes in three different cell lines overexpressing this NR.

### PGA inhibits the differentiation of murine and human CD4^+^ T cells into Th17 cells and inhibits the release of IL-17A from pre-differentiated human Th17 cells

As a next step, it was assessed whether PGA affects murine and human Th17 differentiation. For murine Th17 cells, total splenocytes of OT-II transgenic mice were cultured under Th17 polarizing conditions for three days and simultaneously treated with the vehicle control, SR2211 (1 µM) or PGA at different concentrations. **Figure 6A** shows the workflow while **Figure 6B** depicts the flow cytometric analysis with the percentage of the IL-17A/RORγt double positive population (IL-17^+^/RORγt^+^) after treatment (the full gating strategy is depicted in **Figure S 10**). PGA treatment led to a concentration-dependent decrease of IL-17^+^/RORγt^+^ cells (**Figure 6C**). While PGA significantly reduced IL-17A expression, the overall percentage of RORγt^+^ cells remained over 75 % at all PGA concentrations tested and above 85% at 3 µM PGA and at lower concentrations (**Figure 6D**) except for mouse 1 (**Figure 6D**, black curve) where a decrease in the percentage of RORγ^+^ cells to approximately 50% was observed at 10 µM. This drop coincides with a decrease in cell viability at 10 µM (**Figure S 11**) and could therefore be non-specific, although the percentage of RORγt^+^ (and IL-17A^+^) cells reflect only the viable population (see gating strategy in **Figure S 10**). Thus, this phenomenon could be an experimental artifact.

**Figure 6:**
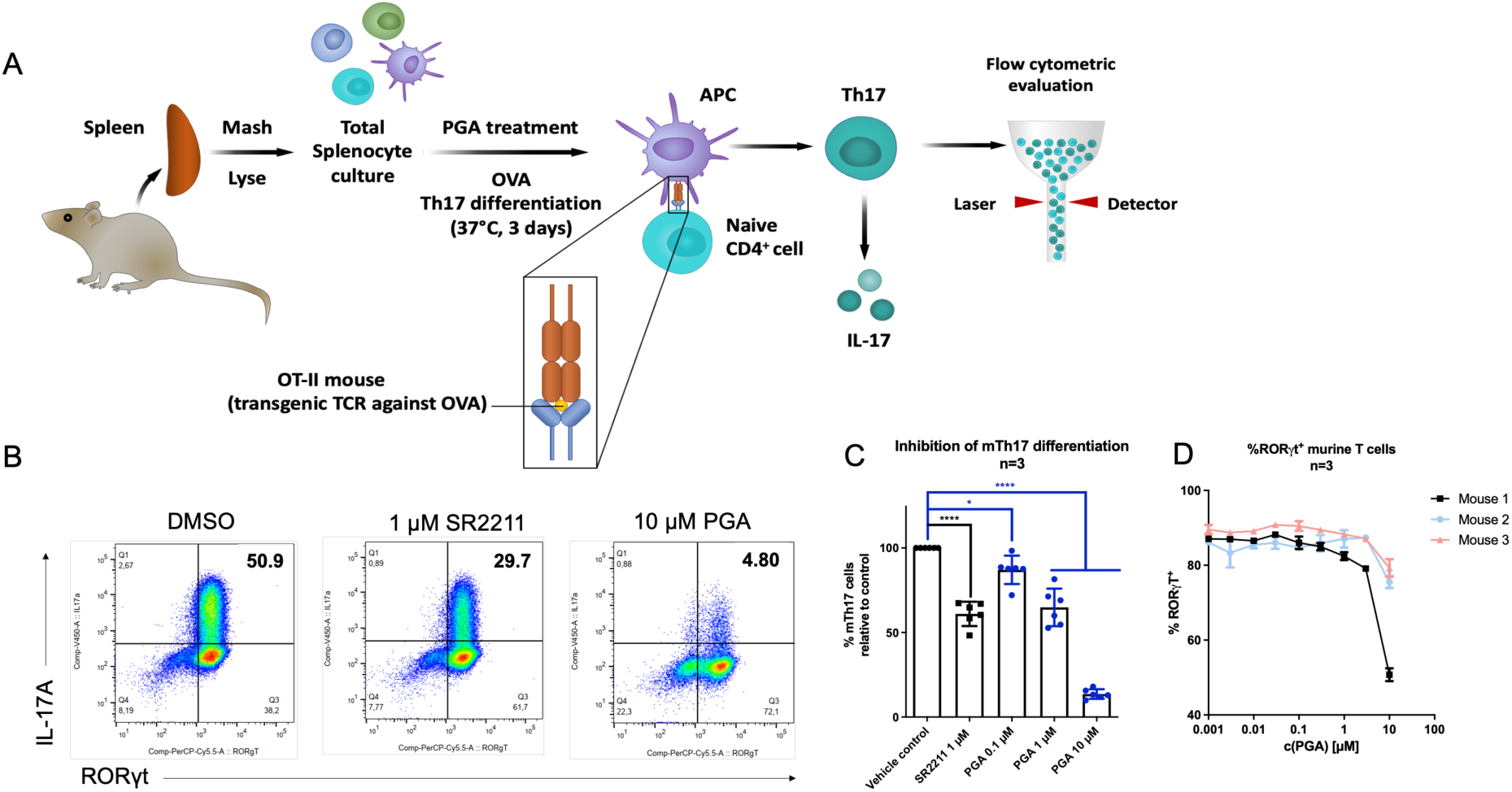
PGA concentration-dependently inhibits murine Th17 differentiation. **A** Schematic overview of the experimental setup measuring murine Th17 differentiation. **B** Flow cytometric analysis of OVA-specific CD4^+^ T-cells activated with OVA peptide and cultured under Th17-polarizing conditions in the presence or absence of PGA for three days. Numbers indicate the percentage of IL-17A^+^/RORγt^+^ cells within viable CD4^+^ T cells. **C** Bar graph showing the percentages of RORγt^+^/IL-17A^+^ CD4^+^ T cells after treatment with SR2211 or PGA relative to vehicle control treatment. Mean ± SD, n=3 in technical duplicates. One-way ANOVA with Dunnett’s test was performed for statistical analysis. **** P ≤ 0.0001, * P ≤ 0.05. **D** The percentage of viable RORγt^+^ cells upon PGA treatment.

Next, naïve human CD4^+^ T cells isolated from peripheral blood mononuclear cells (PBMCs) of healthy anonymous donors were differentiated toward Th17 cells for seven days[35] and simultaneously treated with compounds of interest. **Figure 7A** represents the experimental workflow while **Figure 7B** depicts dot plots of the flow cytometric analysis, which shows that SR2211 (3 µM) and PGA (10 µM) inhibit the differentiation of human Th17 cells compared to vehicle control (the full gating strategy is shown in **Figure S 12**). A concentration-dependent decrease in the percentage of human Th17 cells upon PGA treatment (**Figure 7C**) was observed, confirming the findings obtained from murine cells.

**Figure 7:**
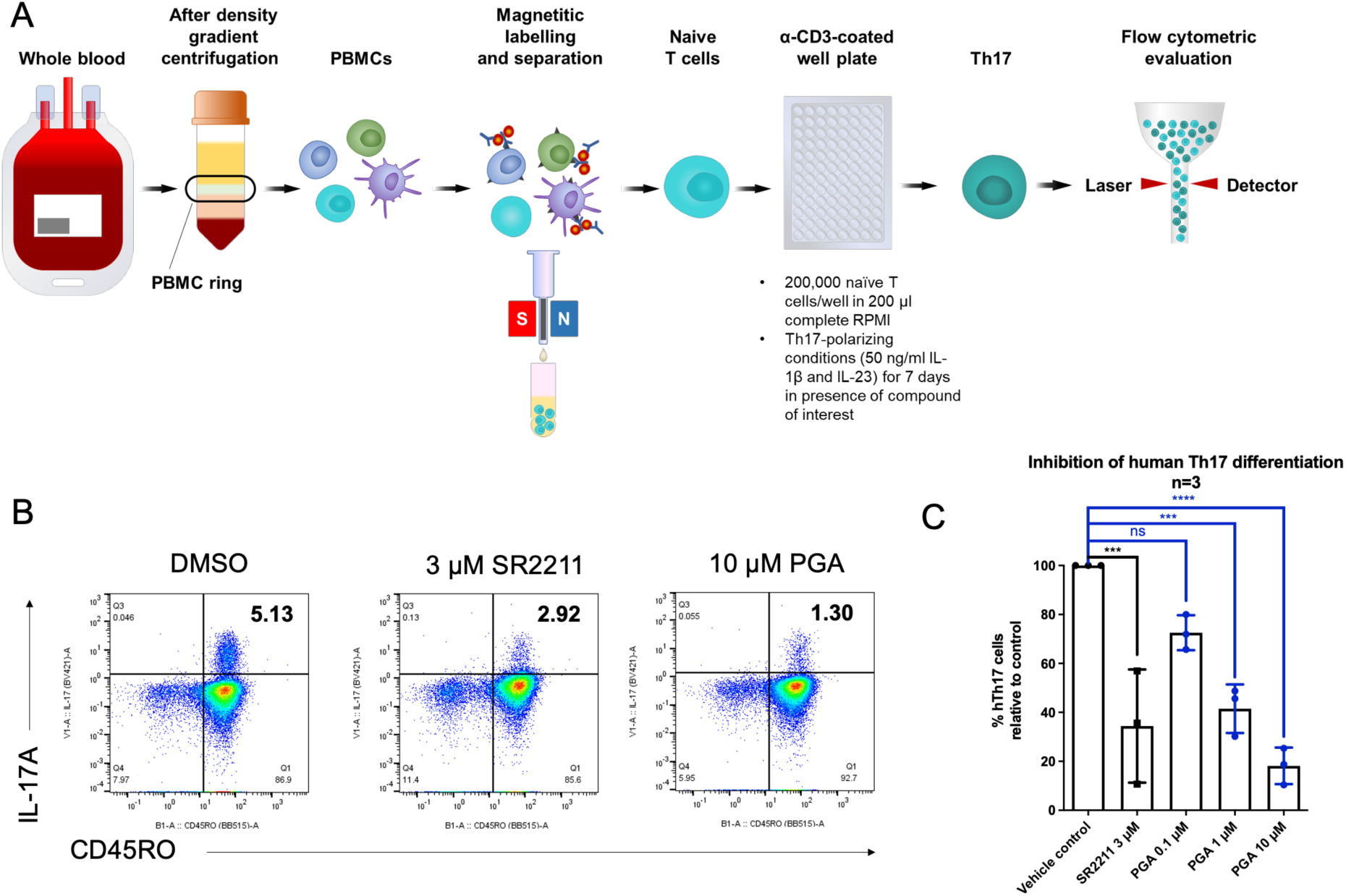
PGA concentration-dependently inhibits human Th17 differentiation. **A** Schematic overview of the experimental setup involving human Th17 differentiation. **B** Flow cytometric analysis of naïve CD4^+^ T-cells activated with plate-bound α-CD3 antibody and cultured under Th17-polarizing conditions in the presence or absence of PGA for seven days. Numbers indicate the percentage of CD45RO^+^ (a marker for IL-17A expressing memory T cells[36]) or CD5^+^/IL-17A^+^ cells. **C** Bar graph showing the percentages of CD45RO^+^ or CD5^+^/IL-17A^+^ helper T cells after treatment with SR2211 or PGA relative to vehicle control treatment. Mean ± SD, n=3 (three healthy, anonymous donors). One-way ANOVA with Dunnett’s test was performed for statistical analysis. **** P ≤ 0.0001, *** P ≤ 0.001.

Finally, the influence of PGA on already differentiated human Th17 cells was explored. To this end, human Th17 cells (CD4^+^CD45RO^+^CXCR3^−^CCR6^+^; **Figure S 13**) were isolated by fluorescence-activated cell sorting (FACS) from enriched CD4^+^ T cells isolated from buffy coats of healthy anonymous donors. Isolated cells were re-stimulated with plate-bound α-CD3 and simultaneously treated with the respective compounds for three days. The workflow is depicted in **Figure 8A**. Dot plots in **Figure 8B** show that SR2211 (3 µM) and PGA (3 µM) reduced the number of IL-17A-producing cells compared to the vehicle control. PGA led to a similar reduction in IL-17A^+^ cells compared to the positive control SR2211 (**Figure 8C**).

**Figure 8:**
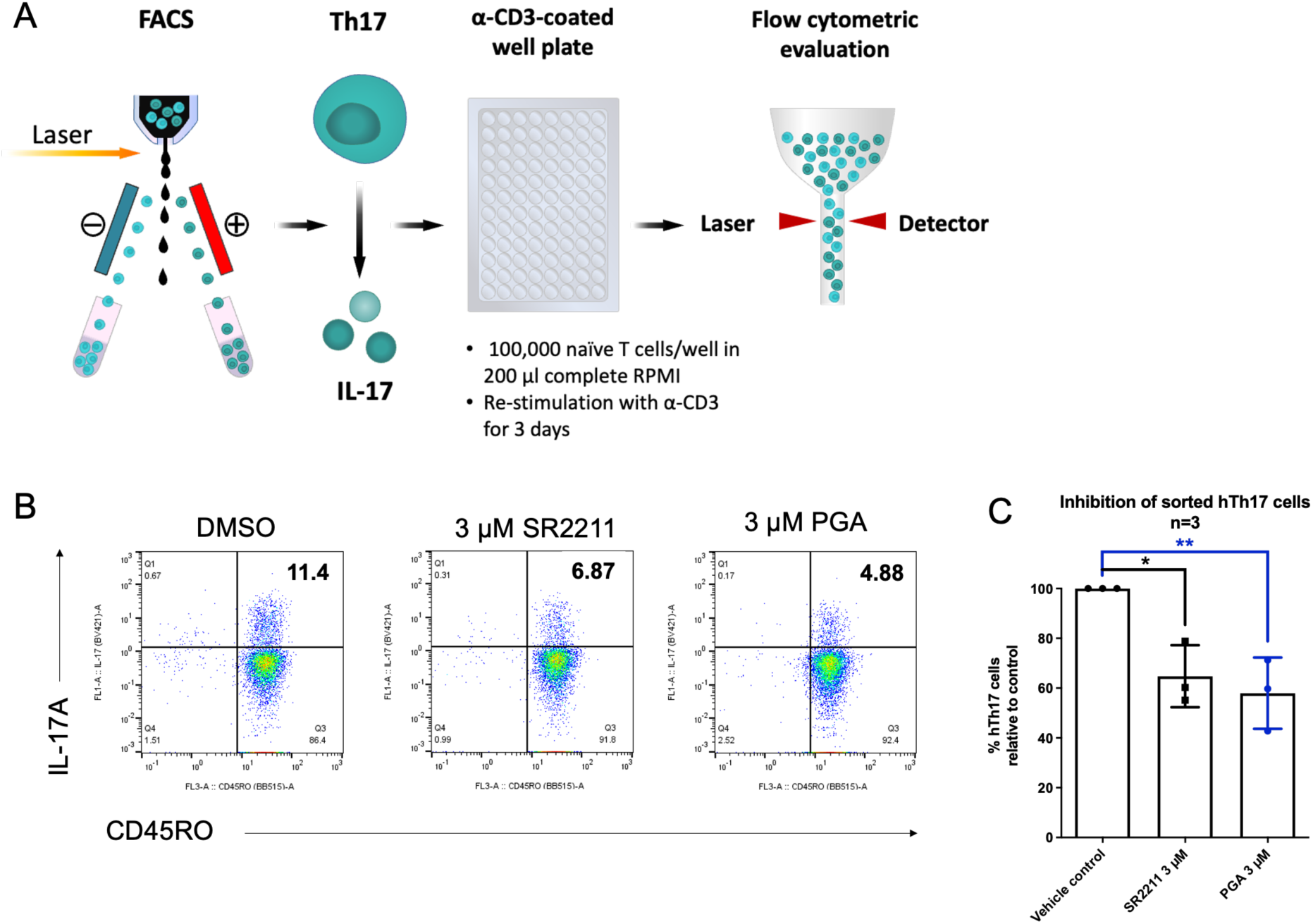
A Schematic overview of the experimental setup using isolated human Th17 cells. **B** Flow cytometric analysis (showing IL-17A and CD45RO) of sorted Th17 cells re-stimulated with plate-bound α-CD3 antibody and cultured in the presence or absence of PGA for three days. Numbers indicate the percentage of cells in the upper right quadrant. **C** Bar graph showing the percentages of CD45RO/IL-17A^+^ CD4^+^ T cells after treatment with SR2211 or PGA relative to vehicle control treatment. Mean ± SD, n=3 (three healthy, anonymous donors). One-way ANOVA with Dunnett’s test was performed for statistical analysis. ** P ≤ 0.01, * P ≤ 0.05.

Collectively, PGA inhibited both murine and human Th17 differentiation in a concentration-dependent manner. Moreover, it was able to decrease the percentage of IL-17A producing cells from already differentiated Th17 cells at 3 µM.

## Discussion

By using a combination of *in vitro*, *in silico*, and *ex vivo* approaches, the oleanane-type triterpenoid PGA, derived from the traditionally used herbal remedy Primulae radix was unveiled as a potent and efficacious inverse agonist of the NR RORγ, the main transcription factor for the differentiation of pro-inflammatory Th17 cells[18]. PGA showed a high potency and efficacy in luciferase assays (IC_50_ approx. 100 nM, I_max_ up to 88%), concentration-dependently reduced RORγ target gene expression in different cell lines and inhibited murine and human Th17 cell differentiation. Additionally, it was able to decrease the percentage of IL-17-producing cells in sorted human Th17 cells, which is consistent with the role of RORγt as a “safeguard” of the Th17 lineage[37].

PGA is a sapogenin obtained by hydrolysis and phytochemical processing of native primula saponins containing the aglycone protoprimulagenin A[28]. The traditional and, to some extent, scientifically validated[10–13] respiratory effects of Primulae radix may at least partly be explained by inhibition of RORγ, a target that has been linked to respiratory disorder treatment as well[38]. This hypothesis is further supported by the high similarity between the saponin profiles of the plant extract and a commercially available cough syrup. However, it remains unclear whether the acidic environment in the stomach is sufficient to facilitate hydrolysis of these saponins and subsequent formation of PGA[39]. This aspect warrants further investigation.

The identified natural product PGA stands out among other known small-molecule inverse agonists of RORγ by its promising potency, suggesting potential therapeutic relevance in the context of Th17-mediated diseases, including psoriasis, psoriatic arthritis, RA, and IBD. Although antibodies against IL-17 like secukinumab are well-established therapeutics for psoriasis and psoriatic arthritis, they surprisingly either show low efficacy or even increase disease severity in RA and IBD, respectively[19]. One explanation could be that Th17 cells change their phenotype from predominantly producing IL-17 to predominantly producing interferon (IFN)-γ in the course of these diseases. Due to a lack of IL-17 production from such ex-Th17 (or nonclassical Th1) cells, anti-IL-17 antibodies are ineffective[40]. Th17 cells have been referred to as “moving targets” due to this plasticity and their ability to alter the cytokines they release[41]. The ensuing therapeutic “loophole” might be effectively closed by employing inhibitors that target RORγt and thus their differentiation rather than the cytokines they secrete. Furthermore, small molecules such as PGA offer advantages such as lower costs and the possibility of oral administration compared to antibodies. The latter point is highly dependent on the pharmacokinetic profile, which is currently unknown for PGA. Given the structural similarity between PGA and OA, it is plausible that PGA (like OA) has low bioavailability [42]. However, semisynthetic derivatives of OA with improved bioavailability have been developed[43], a strategy that could also be applied to PGA. To generate such derivatives, *in silico*-guided semisynthetic modifications of PGA could be utilized. A first step towards this has already been taken in this study i) by confirming direct binding of PGA to the hRORγ LBD and ii) by unveiling the likely binding mode of PGA within the RORγ LBD by molecular docking. However, predictions made by docking still need additional experimental verification, especially since the backbone interaction with F377 using site-directed mutagenesis could not be verified.

The most pressing issue, which potentially limits the therapeutic use of PGA originates from the uncertain safety profile of RORγ inverse agonists. While the findings collected in this study demonstrate non-cytotoxicity of PGA at the used concentrations, the general applicability of RORγ inverse agonists as therapeutics has been questioned. It was shown that *Rorc^−/-^* mice have a high incidence of metastatic thymic lymphomas that tend to metastasize into spleen and liver[44]. The development of lymphoblastic lymphomas was also observed in RORγ adult induced knockout mice[45]. Importantly, treatment of rasH2-Tg hemizygous mice with the RORγ inverse agonist BMS-986251 effectively phenocopied *Rorc*^−/-^ mice by leading to a high incidence of thymic lymphomas and to the discontinuation of this inhibitor[46]. All these findings raise substantial safety concerns when it comes to the therapeutic use of RORγ inverse agonists. In this regard, RORα was suggested as a “safer” therapeutic target compared to RORγ since its inhibition did not cause thymic apoptosis (which was linked to the development of lymphomas[44]) but still suppressed Th17 differentiation and the development and severity of experimental autoimmune encephalomyelitis, a disease model for MS[36]. On the other hand, the RORγ inverse agonist IMU-935 was deemed safe with no dose limiting toxicities in a double blind, placebo-controlled, first-in-human phase 1 study[47]. Prior to that study, authors demonstrated that IMU-935 had no effect on thymocyte maturation *in vitro* and did not completely inhibit RORγ. Instead, it maintained the receptor at approximately 20% of its basal activity in a Gal4-based luciferase assay at the highest concentration used[48]. This suggests that the remaining activity of RORγ is sufficient to support proper thymocyte maturation. Thus, by employing “partial” inverse agonists like IMU-935, detrimental side effects of RORγ inhibition could be avoided[49]. In comparison, 10 µM PGA leaves approximately 30% and 10% of the basal RORγ activity in the Gal4 and full-length luciferase assay, respectively. While this corresponds well to the luciferase data collected for IMU-935, it is ultimately no proof for the safety of PGA.

In summary, PGA, an oleanane-type triterpenoid from the traditionally used herbal medicine Primulae radix was discovered as new RORγ inverse agonist with high potency and efficacy.

## Materials and methods

### Materials

All purchased items including company and identifier can be found in **Table S7**, **Table S8**, and **Table S9**. Gifted plasmids and all used primers are listed in **Table S 10** and **Table S 11**, respectively. All self-designed primers were purchased from Microsynth.

### Compound Stocks, Dilutions, and Identity/Purity Checks

Compounds were dissolved in 100% DMSO under sterile conditions (HERAsafe KS18, Thermo Fisher Scientific). Compound stocks and dilutions were stored at −70 °C until use. For identity and purity checks, these stock solutions were diluted 1:100 with MeOH before analysis.

Compound identity and purity were analyzed using a UHPLC-DAD-CAD-MS system. The setup consisted of an Ultimate 3000 UHPLC system (Thermo Fisher Scientific) coupled to DAD and CAD detectors, using a reversed-phase ACQUITY UPLC CSH C18 column (130 Å, 1.7 μm, 2.1 × 100 mm; Waters Corp.). Mobile phases were A: H_2_O with 0.01% formic acid, and B: acetonitrile; both were degassed prior to use. A 10-minute binary gradient was applied at a flow rate of 350 μl/min: 0–1 min, 80% B; 2–6 min, 80–98% B; 6–8 min, 98% B; 8–10 min, re-equilibration with 80% B. Samples (10 μl, in MeOH) were injected, followed by a blank to ensure column cleaning and re-equilibration. Purity was assessed via DAD and CAD chromatograms. Compound identity was confirmed by mass spectrometry using an LTQ-XL linear ion trap mass spectrometer (Thermo Fisher Scientific) with a HESI source (capillary temp: 350 °C; sheath/aux/sweep gas: 54/12/3 arbitrary units; spray voltage: 3.5 kV). Data were acquired in both positive and negative ion modes over an m/z range of 100–2000 Dalton.

### Cell culture

Cell culture techniques were performed under sterile conditions. All incubation steps were performed at 37 °C and 5% CO_2_. HEK293 and EL-4 (mRORγt) cells were cultured in DMEM, supplemented with 10% fetal bovine serum (FBS), 2 mM L-glutamine, and Penicillin-Streptomycin (100 U/ml penicillin and 100 µg/ml streptomycin) (“complete DMEM”). Jurkat T and HepG2 cells were cultured in RPMI 1640 and EMEM containing the same supplements, respectively. All media contained phenol red. Cells were subcultured every 2-3 days and only used up to in-house passage number of 30. Cell counting and viability were checked using a Vi-CELL XR Cell Viability Analyzer (Beckman Coulter). Where indicated, 5% charcoal-stripped FBS replaced 10% FBS (“stripped” media). Where indicated, stripped DMEM without phenol red was used.

### Luciferase reporter gene assay

8 × 10^6^ HEK293 cells in complete DMEM were seeded onto a 150 mm dish and incubated for 5 hours. Afterwards, cells were transfected with 5 µg of a plasmid encoding the NR, 5 µg of a plasmid encoding the luciferase reporter including the respective response element, and 3 µg of a plasmid encoding eGFP using the calcium phosphate co-precipitation method[50]. After overnight incubation, medium was changed to complete DMEM. After 4-5 hours, cells were trypsinized (0.05% trypsin + 537 µM Na_2_-EDTA*2H_2_O in 1000 ml PBS), resuspended in stripped DMEM and seeded into a 96-well plate at a density of 5 × 10^4^ cells per well. Cells were treated with the vehicle control, the positive control, or the compounds of interest and incubated for 18 hours. Afterwards, cells were checked microscopically for signs of toxicity or compound precipitation. Medium was removed and cells were frozen at −70 °C for at least 1 hour. Cells were lysed with the Reporter Lysis 5X Buffer enriched with coenzyme A (CoA; 450 µM) and dithiothreitol (DTT; 5 mM). RFU and RLU values were measured on a Tecan Spark (Tecan Group), the latter after applying 50 µl adenosine triphosphate (ATP) and D-luciferin *via* injectors A and B, respectively. The RLU/RFU ratio was calculated to account for differences in cell number and transfection efficiency. RLU/RFU values were then normalized to the vehicle control and are expressed as “fold activation”.

### General experimental procedures connected to the isolation of PGA

Normal phase flash chromatography was performed on an Interchim puriFlash 4250 system, equipped with an evaporative light scattering detector (ELSD), a photodiode array (PDA) and a fraction collector, controlled by Interchim Software. A PuriFlash Silica HP column (15 µm, 120 g) served as stationary phase and the mobile phase consisted of *n*-hexane, acetone, and methanol. Semi-preparative supercritical fluid chromatography (SFC) was performed on a Waters Prep-15 System equipped with an ELSD, a PDA, and a fraction collector. A Waters Viridis Prep BEH column (5 µm; 10 × 250 mm) served as the stationary phase and data were analyzed using MassLynx (Waters Corp.). The mobile phase consisted of a supercritical CO_2_/organic modifier (MeOH) gradient (temperature, 45 °C; flow rate, 15 ml/min). The fractions obtained from all chromatographic steps were analyzed by TLC (mobile phase: CH_2_Cl_2_-acetone, 5.5:1); stationary phase: Merck silica gel 60 PF_254_, detected after derivatization with vanillin/H_2_SO_4_ (5% in MeOH) under both visible light and UV_254_ and UV_366_. Ultra-high-performance supercritical fluid chromatography (UHPSFC) was carried out using an Acquity UPC^2^ (ultra-performance convergence chromatography) instrument comprising a sample-, binary solvent-, column-, isocratic solvent- and convergence manager with a PDA detector and a Quadrupole Dalton (QDa) MS detector equipped with electrospray ionization (ESI) in the positive and negative modes: capillary voltage, 0.8 kV; nebulizer, 0.4 bar (N_2_); dry gas flow, 4 l/min (N_2_); probe temperature, 500 °C. The instrument was controlled *via* Empower 3 software (Waters Corp.).

1D and 2D NMR experiments were performed by using a Bruker Avance 500 NMR spectrometer (UltraShield) with a 5 mm switchable probe (TCI Prodigy CryoProbe, 5 mm, triple resonance inverse detection probe head) with z axis gradients and automatic tuning and matching accessory (Bruker BioSpin). The sample (~2 mg) were measured at 298 K in fully deuterated CDCl_3_ referenced to the residual non-deuterated solvent signals. The resonance frequency for ^1^H-NMR was 500.13 MHz and for ^13^C-NMR 125.75 MHz. Standard 1D and gradient-enhanced (ge) 2D experiments, like HSQC, HMBC, and NOESY were used as supplied by the manufacturer.

### Extraction, isolation, and identification of PGA

The dried cut roots of *Primula* sp. (~1 kg) were macerated with 3 L MeOH (at 22°C, three times for 48 hours each). After removal of the solvent under vacuum, the extract was reconstituted in a mixture of MeOH and water (1:1) and separated *via* liquid-liquid partition using petroleum ether. The combined aqueous MeOH phases were then acidified by adding concentrated HCl to reach a pH value of 1.5. To hydrolyze the contained saponins, this mixture was stirred and heated at 95°C for several hours. Successful hydrolysis of saponins was confirmed via TLC. The obtained extract (PRA) was partitioned by CH_2_Cl_2_ to separate aglycones (PRA_A), i.e. sapogenins (in the CH_2_Cl_2_ phase) from sugars (PRA_S) in the remaining aqueous phase. The PRA_A fraction (13 g) was then subjected to normal phase flash chromatography resulting in eight fractions (1.1 to 1.8). Fraction 1.4 (890 mg) was further separated using Sephadex LH 20 column chromatography (mobile phase: CH_2_Cl_2_-acetone, 85:15) to give four fractions (2.1 to 2.4). Fraction 2.4 (270 mg) was then subjected to semi-preparative SFC resulting in seven fractions (3.1 to 3.7). Fraction 3.5 was identified as primulagenin A (PGA, 20 mg). The purity of PGA was determined to be 97.4% by ^1^H-NMR [51].

### Organic synthesis of triterpenoids

#### General information

All reactions requiring anhydrous conditions were performed under argon atmosphere in oven dried glassware applying *Schlenk* technique. Reactions carried out at 0 °C employed an ice bath. All commercially purchased chemicals were used without further purification unless otherwise noted. THF was dried by storing it over 4 Å molecular sieves under argon atmosphere. Infrared spectra were obtained from neat solids or liquids on a Bruker Alpha with a Platinum-ATR unit. Melting points were determined in open capillary tubes on a Cole Parmer Stuart SMP11 melting point apparatus and are uncorrected. Optical rotations were measured using an Anton Paar MCP 100 polarimeter. Concentrations (c) of specific rotations are given in g/100 ml. High resolution mass spectra (HRMS) were measured with a Bruker maXis HD ESI-QTOF. NMR spectra were recorded on a Bruker AVANCE III ASCEND 400 MHz spectrometer with BBFO-PLUS probe or on Bruker AVANCE III HD Ultrashield 500 MHz spectrometer with TCI H/F-C-N Prodigy Kryo-probe head at 298 K. As internal standard, the residual signal of deuterated chloroform 7.26/77.16 ppm (*δ* ^1^H/^13^C ppm) was used[52]. All relevant signals are listed as follows: chemical shift, multiplicity (s = singlet, d = doublet, dd = doublet of doublets, ddd = doublet of doublet of doublets, t = triplet, td = triplet of doublets, m = multiplet), coupling constant(s) and number of protons/carbons. Reactions were monitored by thin layer chromatography (TLC) on silica-coated aluminium plates (Macherey-Nagel, DC Kieselgel Alugram^®^ Xtra SIL G/UV254, layer thickness 0.2 mm). Visualization was performed with vanillin staining (6.0 g vanillin, 1.5 ml sulfuric acid >95%, 95 ml ethanol). Flash column chromatography was performed using silica 60 (particle size 0.040-0.063 nm, 230-400 mesh, Macherey-Nagel) at room temperature using a MPLC-system (Biotage^®^ Selekt).

#### Synthesis of primulagenin A

Under argon-atmosphere, lithium aluminium hydride LiAlH_4_ (powder, 76.3 mg, 2.01 mmol, 3.80 equiv.) was added in portions at 0 °C to a solution of echinocystic acid (250 mg, 529 µmol, 1.00 equiv.) in anhydrous THF (60 ml). The reaction mixture was heated under reflux for 3 hours, cooled to room temperature and stirred at this temperature for 18 hours. Afterwards, the reaction mixture was cooled to 0 °C, quenched by the successive addition of water (10 ml), 4 M aqueous NaOH (10 ml) and water (10 ml) and warmed to room temperature within 15 minutes. Subsequently, the reaction mixture was diluted with methyl *tert*-butyl ether (MTBE) (30 ml) and washed with sat. aqueous NaHCO_3_-solution (30 ml), followed by the extraction of the aqueous phase with MTBE (3 × 30 ml). The combined organic phases were washed with brine (60 ml), dried over MgSO_4_, filtered and the solvent was removed *in vacuo*. The obtained residue was purified by column chromatography (SiO_2_, CH_2_Cl_2_/MeCN 9:1 → 5:1) to yield primulagenin A (210 mg, 457 µmol, 86%) as white solid.

**TLC**: R*_f_* = 0.30 (CH_2_Cl_2_/MeCN 6:1).

**Mp.**: 234 °C.

[α]^20^_D_: +24.0 (c = 0.10, CHCl_3_).

**IR** (ATR): *ṽ* (cm^−1^) = 3888, 2934, 2157, 1457, 1382, 1337, 1301, 1258, 1192, 1080, 1038, 998,

727, 656, 631.

**^1^H-NMR** (400 MHz, CDCl_3_): *δ* (ppm) = 0.73─0.77 (m, 1 H, 5-H), 0.79 (s, 3 H, 24-H_3_), 0.92 (s,

6 H, 29-H_3_, 30-H_3_), 0.93 (s, 3 H, 26-H_3_), 0.94 (s, 3 H, 25-H_3_), 0.95─0.99 (m, 1 H, 1-H_A_), 1.00 (s, 3 H, 23-H_3_), 1.08 (ddd, *J* = 1.2, 3.8, 12.4 Hz, 1 H, 19-H_A_), 1.24─1.28 (m, 1 H, 21-H_A_), 1.34 (s, 3 H, 27-H_3_), 1.34─1.40 (m, 1 H, 15-H_A_), 1.38─1.43 (m, 2 H, 6-H_A_, 7-H_A_), 1.52─1.56 (m, 1 H, 7-H_B_), 1.54─1.60 (m, 2 H, 6-H_B_, 9-H), 1.57─1.62 (m, 1 H, 21-H_B_), 1.59─1.64 (m, 4 H, 2-H_2_, 22-H_2_), 1.62─1.66 (m, 1 H, 1-H_B_), 1.86─1.90 (m, 2 H, 11-H_2_), 1.89─1.94 (m, 1 H, 15-H_B_), 1.91─1.96 (m, 1 H, 18-H), 2.04─2.10 (m, 1 H, 19-H_B_), 3.22 (dd, *J* = 4.7, 11.1 Hz, 1 H, 3-H), 3.32 (s, 2 H, 28-H_2_), 4.05 (t, *J* = 4.8 Hz, 1 H, 16-H), 5.32 (t, *J* = 3.7 Hz, 1 H, 12-H).

**^13^C-NMR** (126 MHz, CDCl_3_): *δ* (ppm) = 15.7 (C-24), 15.9 (C-25), 17.3 (C-26), 18.4 (C-6), 23.5 (C-11), 25.6 (C-30), 26.4 (C-22), 27.4 (C-2), 27.4 (C-27), 28.2 (C-23), 30.5 (C-20), 32.9 (C-7), 32.9 (C-29), 34.9 (C-15), 35.4 (C-21), 37.1 (C-10), 38.8 (C-1), 38.9 (C-4), 40.1 (C-8), 40.7 (C-17), 41.7 (C-14), 42.8 (C-18), 47.1 (C-9), 47.1 (C-19), 55.4 (C-5), 70.9 (C-28), 75.1 (C-16), 79.1 (C-3), 122.9 (C-12), 143.1 (C-13).

**HRMS** (ESI^+^) = *m/z* calcd: 481.3652 C_30_H_50_NaO_3+_; found: 481.3650.

**C_30_H_50_O_3_** (458.72 g/mol).

#### Synthesis of erythrodiol

Under argon-atmosphere, lithium aluminium hydride (LiAlH_4_) (powder, 190 mg, 4.99 mmol, 3.80 equiv.) was added in portions at 0 °C to a solution of oleanolic acid (600 mg, 1.31 mmol, 1.00 equiv.) in anhydrous THF (30 ml). The reaction mixture was heated to reflux for 3 hours, cooled to room temperature and stirred at this temperature for 16 hours. Afterwards, the reaction mixture was cooled to 0 °C, quenched by the successive addition of water (10 ml), 4 M aqueous NaOH (10 ml) and water (10 ml) and warmed to room temperature within 15 minutes. Subsequently, the reaction mixture was diluted with methyl *tert*-butyl ether (MTBE) (40 ml) and was washed with water (2 × 30 ml), followed by the extraction of the aqueous phase with MTBE (3 × 30 ml). The combined organic phases were washed with brine (50 ml), dried over MgSO_4_, filtered and the solvent was removed *in vacuo*. The obtained residue was purified by column chromatography (SiO_2_, PE/EtOAc 19:1 → 3:1) to yield erythrodiol (400 mg, 903 µmol, 69%) as white solid.

**TLC**: R*_f_* = 0.34 (PE/EtOAc 4:1).

**Mp.**: 224 °C.

[α]^20^_D_: +72.5 (c = 0.10, CHCl_3_).

**IR** (ATR): *ṽ* (cm^−1^) = 3327, 2924, 2867, 1462, 1365, 1305, 1189, 1148, 1093, 1041, 1000, 816,

716, 603, 521.

**^1^H-NMR** (400 MHz, CDCl_3_): *δ* (ppm) = 0.72─0.75 (m, 1 H, 5-H), 0.79 (s, 3 H, 24-H_3_), 0.87 (s, 3 H, 30-H_3_), 0.89 (s, 3 H, 29-H_3_), 0.93 (s, 3 H, 25-H_3_), 0.94 (s, 3 H, 26-H_3_), 0.94─0.99 (m, 1 H, 1-H_A_), 0.97─1.02 (m, 1 H, 15-H_A_), 1.00 (s, 3 H, 23-H_3_), 1.04─1.08 (m, 1 H, 19-H_A_), 1.14─1.18 (m, 1 H, 16-H_A_), 1.17 (s, 3 H, 27-H_3_), 1.17─1.21 (m, 1 H, 21-H_A_), 1.27─1.33 (m, 1 H, 21-H_B_), 1.31─1.35 (m, 1 H, 7-H_A_), 1.32─1.37 (m, 1 H, 22-H_A_), 1.37─1.43 (m, 1 H, 6-H_A_), 1.49─1.55 (m, 2 H, 7-H_B_, 22-H_B_), 1.52─1.58 (m, 2 H, 6-H_B_, 9-H), 1.56─1.63 (m, 2 H, 2-H_2_), 1.60─1.64 (m, 1 H, 1-H_B_), 1.67─1.73 (m, 1 H, 15-H_B_), 1.70─1.75 (m, 1 H, 19-H_B_), 1.83─1.90 (m, 2 H, 11-H_2_), 1.86─1.93 (m, 1 H, 16-H_B_), 1.96─2.00 (m, 1 H, 18-H), 3.21─3.23 (m, 2 H, 28-H_A_, 3-H), 3.55 (d, *J* = 10.9 Hz, 1 H, 28-H_B_), 5.19 (t, *J* = 3.6 Hz, 1 H, 12-H).

**^13^C-NMR** (126 MHz, CDCl_3_): *δ* (ppm) = 15.7 (C-25), 15.7 (C-24), 16.9 (C-26), 18.5 (C-6), 22.1 (C-16), 23.7 (C-11), 23.7 (C-30), 25.7 (C-15), 26.1 (C-27), 27.4 (C-2), 28.2 (C-23), 31.1 (C-20), 31.2 (C-22), 32.7 (C-7), 33.3 (C-29), 34.2 (C-21), 37.1 (2 C, C-10, C-17), 38.7 (C-1), 38.9 (C-4), 39.9 (C-8), 41.9 (C-14), 42.5 (C-18), 46.6 (C-19), 47.7 (C-9), 55.3 (C-5), 69.8 (C-28), 79.1 (C-3), 122.5 (C-12), 144.3 (C-13).

**HRMS** (ESI^+^) = *m/z* calcd: 465.3703 C_30_H_50_NaO_2+_ found: 465.3717.

**C_30_H_50_O_2_** (442.73 g/mol).

#### Synthesis of oleanolic aldehyde

Under argon-atmosphere, erythrodiol (375 mg, 847 µmol, 1.00 equiv.) was dissolved in anhydrous CH_2_Cl_2_ (12 ml). Then, 2,2,6,6-tetramethylpiperidinyloxyl (TEMPO) (26.5 mg, 169 µmol, 0.20 equiv.) and bis(acetoxy)iodobenzene (BAIB) (409 mg, 1.27 mmol, 1.50 equiv.) were added to the reaction mixture and the mixture was stirred at RT for 16 hours. Upon reaction control (TLC), another portion of BAIB (81.1 mg, 254 µmol, 0.30 equiv.) and TEMPO (26.5 mg, 169 µmol, 0.20 equiv.) were added, and the reaction mixture was stirred at RT for 3 hours and additionally at 40 °C for 2 hours. The reaction mixture was quenched with sat. aqueous NaHSO_3_-solution (5 ml) and the organic phase was washed with sat. aqueous NaHCO_3_-solution (10 ml). The aqueous phase was extracted with CH_2_Cl_2_ (3 × 10 ml) and the combined organic phases were washed with brine (25 ml), dried over MgSO_4_, filtered and the solvent was removed *in vacuo*. The obtained residue was purified by column chromatography (SiO_2_, PE/EtOAc 19:1 → 6:1) yielding oleanolic aldehyde (160 mg, 362 µmol, 43%) as white solid.

**TLC**: R*_f_* = 0.52 (PE/EtOAc 4:1).

**Mp.**: 176 °C.

[α]^20^_D_: +102.0 (c = 0.10, CHCl_3_).

**IR** (ATR): *ṽ* (cm^−1^) = 3513, 2934, 2859, 1712, 1460, 1367, 1294, 1177, 1141, 1096, 1043, 998,

921, 752, 601.

**^1^H-NMR** (400 MHz, CDCl_3_): *δ* (ppm) = 0.70─0.74 (m, 1 H, 5-H), 0.74 (s, 3 H, 26-H_3_), 0.78 (s, 3 H, 24-H_3_), 0.91 (s, 3 H, 25-H_3_), 0.91 (s, 3 H, 30-H_3_), 0.92 (s, 3 H, 29-H_3_), 0.95─0.99 (m, 1 H, 1-H_A_), 0.99 (s, 3 H, 23-H_3_), 1.08 (ddd, *J* = 2.5, 4.2, 13.5 Hz, 1 H, 15-H_A_), 1.14 (s, 3 H, 27-H_3_), 1.17─1.22 (m, 2 H, 22-H_A_, 19-H_A_), 1.24─1.33 (m, 3 H, 21-H_2_, 7-H_A_), 1.35─1.39 (m, 1 H, 6-H_A_), 1.42─1.48 (m, 2 H, 7-H_B_, 22-H_B_), 1.49─1.54 (m, 1 H, 9-H), 1.51─1.55 (m, 1 H, 6-H_B_), 1.55─1.58 (m, 1 H, 16-H_A_), 1.57─1.63 (m, 3 H, 2-H_2_, 1-H_B_), 1.62─1.67 (m, 1 H, 15-H_B_), 1.66─1.70 (m, 1 H, 19-H_B_), 1.85─1.90 (m, 2 H, 11-H_2_), 1.98 (td, *J* = 4.2, 13.5 Hz, 1 H, 16-H_B_), 2.63 (dd, *J* = 4.4, 13.8 Hz, 1 H, 18-H), 3.20─3.22 (m, 1 H, 3-H), 5.34 (t, *J* = 3.7 Hz, 1 H, 12-H), 9.40 (s, 1 H, 28-H).

**^13^C-NMR** (126 MHz, CDCl_3_): *δ* (ppm) = 15.5 (C-25), 15.7 (C-24), 17.2 (C-26), 18.4 (C-6), 22.2 (C-16), 23.6 (C-11), 23.6 (C-30), 25.7 (C-27), 26.9 (C-15), 27.3 (C-2), 27.9 (C-22), 28.3 (C-23), 30.8 (C-20), 32.9 (C-7), 33.2 (C-29), 33.3 (C-21), 37.1 (C-10), 38.6 (C-1), 38.9 (C-4), 39.7 (C-8), 40.6 (C-18), 41.8 (C-14), 45.7 (C-19), 47.7 (C-9), 49.2 (C-17), 55.3 (C-5), 79.1 (C-3), 123.4 (C-12), 143.1 (C-13), 207.7 (C-28).

**HRMS** (ESI^+^) = *m/z* calcd: 441.3727 C_30_H_49_O_2+_ found: 441.3753.

**C_30_H_48_O_2_** (440.71 g/mol).

### Resazurin conversion assay (cell viability assay)

For resazurin conversion assays, the different cell lines were treated with the vehicle control, the cytotoxic positive control (digitonin at 20 or 50 µg/ml), or the compounds of interest for different periods of time (in accordance with the treatment duration in the respective assay). Afterwards, the medium was carefully removed and replaced with stripped DMEM medium without phenol red containing 10 µg/ml resazurin sodium salt. Cells were incubated for 5 hours and then measured on a Tecan Spark at λ_em_ = 590 nm. Cell viability was calculated as follows: 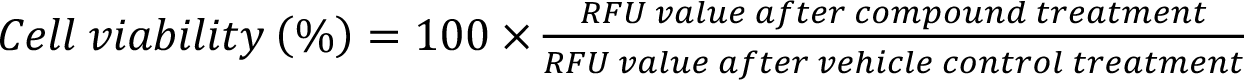. Cytotoxicity was defined as cell viability < 70% according to the ISO 10993-5 guidelines[53].

### Thermal stability assay

PGA and T0901317 were dissolved in absolute ethanol at a concentration of 10^−2^ M. Compounds were stored at −20 °C until use.

The sequence encoding the His-hRORγ (264-518) LBD was inserted into a pET15b expression plasmid. The protein was expressed in *Escherichia coli* BL21 DE3 by induction with 1 mM IPTG at an OD600 of ~0.8 and incubation at 22°C for 150 minutes. Soluble proteins were purified on Ni Hitrap FFcrude column (Cytiva), followed by His tag removal by thrombin cleavage and by size exclusion chromatography on HiLoad Superdex200 column (Cytiva) equilibrated in 20mM Tris-HCl pH7.5, 150mM NaCl, 1mM TCEP. Purity and homogeneity of the hRORγ LBD was assessed by SDS-PAGE. The purified protein was concentrated to 0.5-1.0 mg/ml with an Amicon Ultra 10 kDa MWCO. The aliquots of purified LBD were incubated with 4 equivalent ligands or vehicle for thermal stability experiment.

Fluorescence based thermal experiments were performed using Prometheus NT.48 (NanoTemper Technologies) with grade standard capillaries containing 10 μl RORγ LBD with different ligands. The temperature was increased by a rate of 1 °C/min from 20 to 95 °C and the fluorescence at emission wavelengths of 330 nm and 350 nm was measured for 3 technical replicates. NanoTemper PR.Stability Analysis v1.0.2 was used to fit the data and to determine the melting temperatures Tm. Presented data is the average of two biological replicates (one in case of T0901317).

### Molecular docking

An X-ray crystal structure of the RORγ LBD in complex with ursonic acid (PDB structure 6J3N, resolution 1.99 Å) was utilized for docking with Glide[54–56] (part of the Schrödinger Platform, version 2023-4, Schrödinger Inc.). The protein structure was prepared with the Protein Preparation Wizard within the Maestro molecular modeling environment[57] (part of the Schrödinger Platform) using default settings. The preparation included the (i) addition of hydrogen atoms, (ii) assignment of bond orders, (iii) assignment of protonation and metal charge states with Epik[58, 59], (iv) sampling H_2_O orientations and optimization of the hydrogen bond network, and (v) restrained minimization using the OPLS4 force field[60] to converge heavy atoms to an RMSD of 0.30 Å. The 3D structure of PGA and OA were prepared with LigPrep[61] using default settings including assigning ionization states at pH pH 7.4 ± 2.0 with Epik and optimizing geometries with the OPLS4 force field. For docking with Glide, the ligand binding site was defined within the Receptor Grid Generation wizard to dock ligands with similar size to the co-crystallized ligand. Glide Standard Precision (Glide SP) was used for ligand docking and up to 20 docking poses were set for output.

### Site-directed mutagenesis

hRORγ-Gal4 Q286T, H323A, and the double mutant were generated using the QuikChange Lightning Site-Directed Mutagenesis Kit according to the manufacturer’s instructions. In all cases, 10 ng template (i.e., hRORγ-Gal4 for generation of the Q286T and H323A mutants or hRORγ-Gal4 H323A for generation of the double mutant) were PCR-amplified using the mutagenic forward and reverse primers listed **Table S 11**. All mutagenic primers were designed using the QuikChange Primer Design tool (https://www.agilent.com/store/primerDesignProgram.jsp). PCR settings are shown in **Table S 3**. After performing the PCR reaction, the PCR product was DpnI digested at 37 °C for 5 minutes and transformed into XL1-Blue competent *E. coli*[62]. After overnight incubation at 37°C, a single colony was picked and added to 5 ml of lysogeny broth (LB) medium containing 50 µg/ml zeocin (“pre-culture”). After shaking for 5 hours at 37 °C, the pre-culture was added to 400 ml of LB medium containing 50 µg/ml zeocin and shaken at 37 °C overnight. After centrifugation of the main culture at 5691 × g for 10 minutes and removal of the supernatant, plasmid preparation was done using the PureLink HiPure Plasmid Midiprep Kit according to the manufacturer’s instructions. Plasmid purities and yields were checked using a NanoDrop 2000c (Thermo Fisher Scientific). hRORγ-Gal4 mutants were checked for the presence of the right mutations by Sanger sequencing (Microsynth) using the sequencing primers listed in **Table S 11**.

### Cloning of a mRORγt-pIRES2-eGFP Vector

First, mRORγt was PCR-amplified out of the full-length murine RORγt vector using the Herculase II Fusion DNA Polymerase kit according to the manufacturer’s instructions. For PCR-amplification, primers including a NheI (mRORgt pIRES2-eGFP NheI fwd) and a XhoI (mRORgt pIRES 2-eGFP XhoI rev) restriction site to the 5’ and 3’ end of the fragment, respectively, were used. Primers were designed using SnapGene Viewer (Dotmatics) and their sequences are listed in **Table S 11**. PCR conditions are shown in **Table S 4**. The PCR product (“fragment”) was separated on a 1% agarose gel containing 0.5X SYBR Safe and extracted using the Monarch DNA Gel Extraction Kit according to the manufacturer’s instructions. After double digestion of the fragment and vector (pIRES2-eGFP) with NheI-HF and XhoI[63], separation on a 1% agarose gel, and DNA gel extraction, ligation with a 7x molar excess of fragment relative to vector was performed using the T4 DNA ligase[64]. Transformation and plasmid preparation were done as described in the **“Site-directed mutagenesis”** section with minor adaptions: 5 µl (instead of 2 µl) ligation product was transformed, and a different selection antibiotic (ampicillin, 50 µg/ml) was used. Sanger sequencing was employed to confirm the correct insertion of mRORγt within the pIRES2-eGFP vector (primer used: “IRES-for”, see **Table S 11** for its sequence).

### Generation of the stable EL-4-mRORγt cell line

The AseI-linearized[63] mRORγt-pIRES2-eGFP vector was transfected into EL-4 cells using Lipofectamine LTX according to the manufacturer’s instructions. The eGFP^high^ population was sorted twice using a CytoFLEX SRT (Beckman Coulter) and cultured in complete DMEM until confluency was reached each time. Ultimately, a third sorting procedure was performed and a single eGFP^high^ cell was sorted per well in a 96-well plate containing complete DMEM enriched with 1000 µg/ml G418 for selection. The clones were allowed to grow to confluency and were later analyzed using a MACSQuant Analyzer 10 flow cytometer (Miltenyi Biotec) and FlowJo 10.8.1 (BD). All cells were found to be eGFP^+^ compared to WT EL-4 cells (**Figure S 9**). Aliquots of these EL-4-mRORγt cells were frozen and subsequently used for qPCR experiments.

### Quantitative reverse-transcription polymerase chain reaction

All steps prior to qPCR (seeding, treatment, …) for HepG2 and Jurkat T cells were done as described previously[65]. For EL-4-mRORγt cells, 5 × 10^5^ cells were seeded per well of a 24-well plate in 500 µl complete DMEM, treated with compounds for 20-24 hours, and stimulated with phorbol 12-myristate 13-acetate (PMA) and ionomycin using the “Cell Activation Cocktail” (PMA: 40.5 µM, ionomycin: 669.3 µM; 500X) for approx. 4.5 hours. Total RNA was isolated using the innuPREP RNA Mini Kit 2.0 according to the manufacturer’s instructions. 1 µg RNA was reverse-transcribed into cDNA using the High-Capacity cDNA Reverse Transcription Kit according to the manufacturer’s instructions.

qPCR for all cell lines was performed using either the GoTaq Green Master Mix or the Luna Universal qPCR Master Mix on a LightCycler 480 (Roche Diagnostics). All steps were performed according to the manufacturer’s instructions. Measurement setting on the LightCycler 480 and qPCR primers are listed in **Table S 5** (GoTaq) / **Table S 6** (Luna), and **Table S 11**, respectively. qPCR primers were either bought (*G6PC*, *GAPDH*) or self-designed (rest) using Primer-BLAST (https://www.ncbi.nlm.nih.gov/tools/primer-blast/). Primer efficiencies for all self-designed primers were checked and found to be > 95%. Gene expression data were first normalized to the control genes by employing the 2^−ΔΔCt^ method[66] and then normalized to the vehicle control.

### Murine Th17 polarization

Splenocytes were isolated from OT-II transgenic mice (B6.Cg-Tg(TcraTcrb)425Cbn/J, The Jackson Laboratory), and after lysis of erythrocytes with BD Pharm Lyse Lysing Buffer according to the manufacturer’s instructions, stimulated with 4 µg/ml OVA peptide on 48-well plates (2 × 10^6^ cells/well) in 1 ml T cell medium/well RPMI 1640 (supplemented with 10% FCS, GlutaMAX, Penicillin-Streptomycin, and 50 mM β-mercaptoethanol (β-ME)) under Th17 polarizing conditions using 1 ng/ml TGFβ, 20 ng/ml IL-6, 10 ng/ml IL-1β, 10 ng/ml IL-23, 10 µg/ml anti-IL-4, 10 µg/ml anti-IFN-γ for 3 days.

### Extracellular and intracellular staining and flow cytometric analysis of murine Th17 cells

For cytokine analysis, cells were re-stimulated with PMA (1 mg/ml), Ionomycin (1 mg/ml), in the presence of GolgiStop (1:800) and GolgiPlug (1:800). 2 × 10^6^ cells were surface stained using TCR-β (0.2 mg/ml) and CD4 (0.2 mg/ml). Dead cells were excluded using LIVE/DEAD Fixable Aqua Dead Cell Stain Kit according to the manufacturer’s instructions. For intracellular staining, the cells were fixed with Cytofix Fixation Buffer, permeabilized with Perm/Wash Buffer according to the manufacturer’s instructions and stained with the following antibodies: IL-17A (0.2 mg/ml), and RORγt (0.2 mg/ml). Stained cells were measured with a FACSVerse flow cytometer (BD). Data are representative of technical duplicates of three mice that were analyzed in three independent experiments (n=3) and analysed using FlowJo 10.2 software.

### Isolation of human PBMCs and naïve CD4^+^ T cells

Whole blood from healthy, anonymous donors was purchased from the Austrian Red Cross. For PBMC isolation, whole blood was diluted 1:2 with phosphate-buffered saline (PBS; 123 mM NaCl + 10 mM Na_2_HPO_4_ + 3 mM KH_2_PO_4_ in 1000 ml ddH2O, pH 7.4) and then carefully layered upon half of the volume of Lymphopure relative to diluted blood. Density gradient centrifugation was performed at 800 × g for 30 minutes with the break turned off. The PBMC ring was carefully isolated, collected in a tube, topped up to 50 ml with PBS and centrifuged at 800 × g for 5 minutes. After repeating the washing step once, a last washing step was done at 200 × g for 15 minutes. Afterwards, the supernatant was carefully removed, and the pellet resuspended in 15-20 ml separation buffer (0.5% bovine serum albumin (BSA) + 2 mM EDTA in PBS pH 7.2). After counting the PBMCs on the Vi-CELL XR Cell Viability Analyzer, naïve CD4^+^ T cells were isolated using the Naive CD4^+^ T Cell Isolation Kit II (human) according to the manufacturer’s instructions.

### Human Th17 differentiation and flow cytometric analysis

For human Th17 differentiation, 2 × 10^5^ naïve CD4^+^ T cells were seeded per well of a 96-well plate that had been coated with α-CD3 antibody (5 µg/ml) the day before. Cells were polarized using 50 ng/ml IL-1β and 50 ng/ml IL-23 in presence of the vehicle control, the positive control or PGA and incubated for 7 days[35]. For cytokine analysis, cells were re-stimulated with the PMA/ionomycin “Cell Activation Cocktail” (PMA: 40.5 µM, ionomycin: 669.3 µM; 500X) for 2 hours, and with monensin (1000X) for 2 additional hours. Extra- and intracellular staining of Th17 cells was done using the BD Pharmingen Transcription Factor Buffer Set according to the manufacturer’s instructions. Cells were stained with Fixable Viability Stain 780 (live/dead) and antibodies against CD45RO (a marker for IL-17A expressing memory T cells[36]) or CD5, and IL-17A. Flow cytometric measurements were performed on a MACSQuant Analyzer 10 and data was analyzed using FlowJo 10.8.1.

### Sorting and flow cytometric analysis of human Th17 cells

Human CD4^+^ T cells were isolated from buffy coats purchased from the Austrian Red Cross using the EasySep™ Human CD4^+^ T Cell Isolation Kit according to manufacturer’s instructions. Subsequently, cells were stained with the following antibodies: anti-human CD45 Brilliant Violet 421, anti-human CD4 FITC, anti-human CD45RO PerCP-Cy5.5, anti-human CD45RA PE, anti-human CXCR3 PE-Cy7 and anti-human CCR6 Alexa Fluor 647 and Th17 cells were isolated on a FACS AriaII Sorter (BD) according to the CD45^+^CD4^+^CD45RA^−^CD45RO^+^CXCR3^−^CCR6^+^ phenotype (purity > 95% for all donors, see **Figure S 13**). Subsequently, 1 × 10^5^ Th17 cells were seeded per well of a 96-well plate that had been coated with α-CD3 antibody (5 µg/ml) the day before. Cells were treated with the vehicle control, the positive control or PGA and incubated for 3 days. Extra- and intracellular staining and flow cytometric analysis was performed as described in the previous section.

### Statistical analysis

Information about sample sizes, statistical analyses used, and P values determined for the conducted experiments are shown in the figure legends. All statistical analyses were performed using GraphPad prism version 10 (Dotmatics). For comparing two groups, Student’s two-tailed t-tests was performed. For comparing more than two groups, one-way ANOVA with Tukey’s or Dunnett’s post-hoc test was employed. For comparing the best-fit log IC_50_ value of PGA versus another triterpenoid, an extra sum-of-squares F test was performed. Statistical significance was reached below an alpha of 0.05 in all cases.

## Data availability statement

All original data and findings are provided in the article and supplementary material. Further information is available from the corresponding author upon reasonable request.

## Acknowledgements

The authors would like to express their gratitude to Lucia N. Schöberl, Johanna Raab, Christoph Schön, Martin M. Kraus, Raphael Opferkuch, Dominik D. Burkhard, and Romana Blab for their invaluable assistance with the biological assays. The authors thank Marlene Stockinger for her valuable assistance in the isolation of PGA from Primulae radix, Daniel Schachner for his outstanding technical support with the CytoFLEX SRT, and Carole Peluso-Iltis for excellent technical assistance regarding the nanoDSF experiments. The authors would also like to sincerily thank Scarlet Hummelbrunner for her assistance with the human Th17 cell experiments. Moreover, we greatly appreciate Ammar Tahir’s efforts in identity and purity testing of the compounds. Also, we extend our thanks to Andrea Szabo for illustrating Figures 7A, 8A, and 9A. Lastly, authors want to thank Christine Coffey and Lorenza Bertaina for proofreading the manuscript.

## Author contributions

This study was conceptualized by P.F.S., A.F.P., and V.M.D. V.M.D. provided overall supervision and resources for the study. A.F.P. offered mentorship and ongoing guidance. Compounds were synthesized by J.J. under the supervision of N.S., who also provided the resources for the syntheses. PGA was isolated under the supervision of U.G. and J.M.R. Compounds were analyzed by J.J., N.S., and U.G. Biological experiments were performed and analyzed by P.F.S., A.F.P., T.P., and L.B. Experiments regarding nanoDSF were conducted under the supervision of N.R. Computational methods were executed by Y.C. and supervised by J.K. Experiments regarding mouse Th17 cells were conducted by L.B. under the supervision of T.P. and M.B., the latter also provided the resources for these experiments. Experiments regarding sorting of human Th17 cells were conducted by K.G.S., who also provided the resources for these experiments. Visualization was done by P.F.S., T.P., L.B., K.G.S., and Y.C. P.F.S. wrote the original draft of the manuscript, A.F.P. and V.M.D. revised it. Thereafter, all other authors reviewed and edited the manuscript. All authors have read and agreed to the published version of the manuscript.

