## Supporting Information for "Primulagenin A is a potent inverse agonist of the nuclear receptor RAR-related orphan receptor gamma (RORγ)"

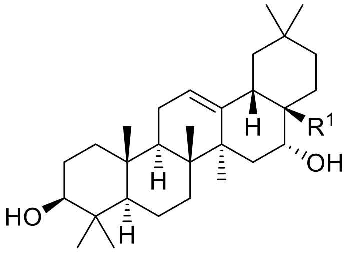


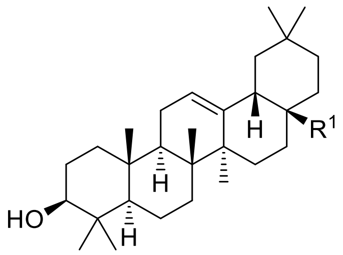


**Echinocystic acid (EA)** $R^1$ = COOH
**Primulagenin D (PGD)** $R^1$ = CHO
**Primulagenin A (PGA)** $R^1$ = $CH_2OH$

**Oleanolic acid (OA)** $R^1$ = COOH
**Oleanolic aldehyde (OAL)** $R^1$ = CHO
**Erythrodiol (ED)** $R^1$ = $CH_2OH$
**β-Amyrin (β-Amy)** $R^1$ = $CH_3$

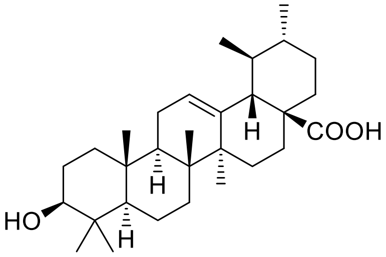


**Ursolic acid (UA)**

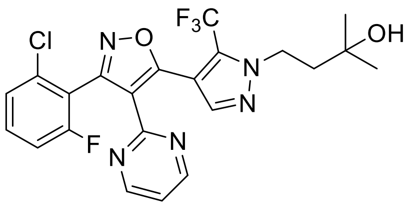


**IMU-935**

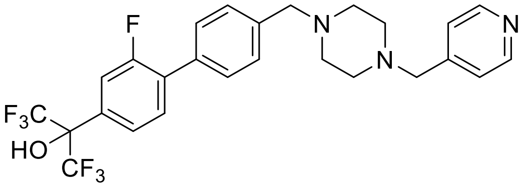


**SR2211**

***Figure S 1:*** *Structural overview of all compounds discussed in this work.*

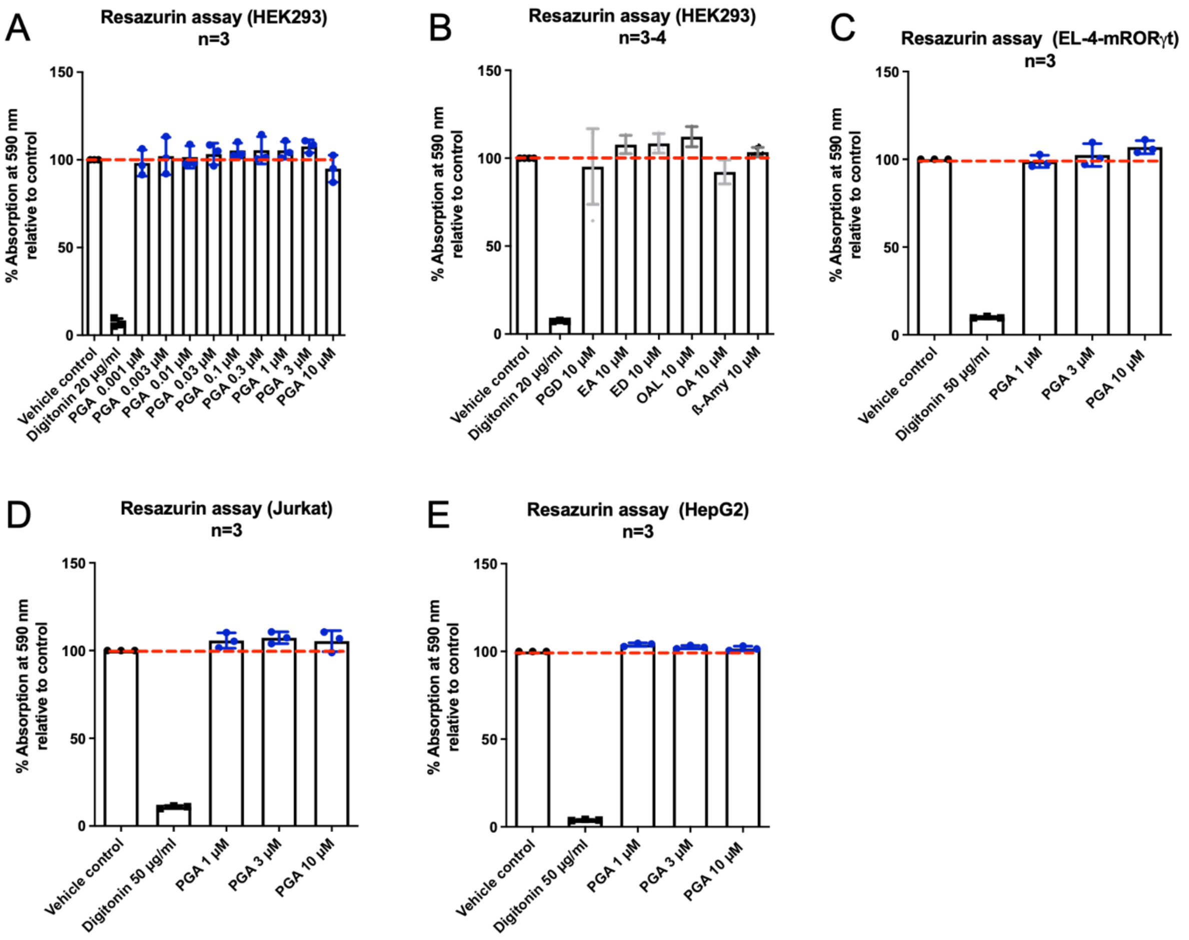


***Figure S 2: PGA and structurally related triterpenoids show not cytotoxicity in any of the cell lines used for other assays. A-E*** *Cells were treated with PGA at the indicated concentrations and were then incubated for the respective durations corresponding to the assays in which they were used. After adding resazurin sodium salt (10 µg/ml), cells were incubated, RFU values were measured at 590 nm and normalized to the vehicle control. Mean ± SD, n=3 in technical quadruplicates (except for the vehicle control, digitonin 50 µg/ml, and PGD at 10 µM in panel* ***B*** *– n=4 in technical quadruplicates).*

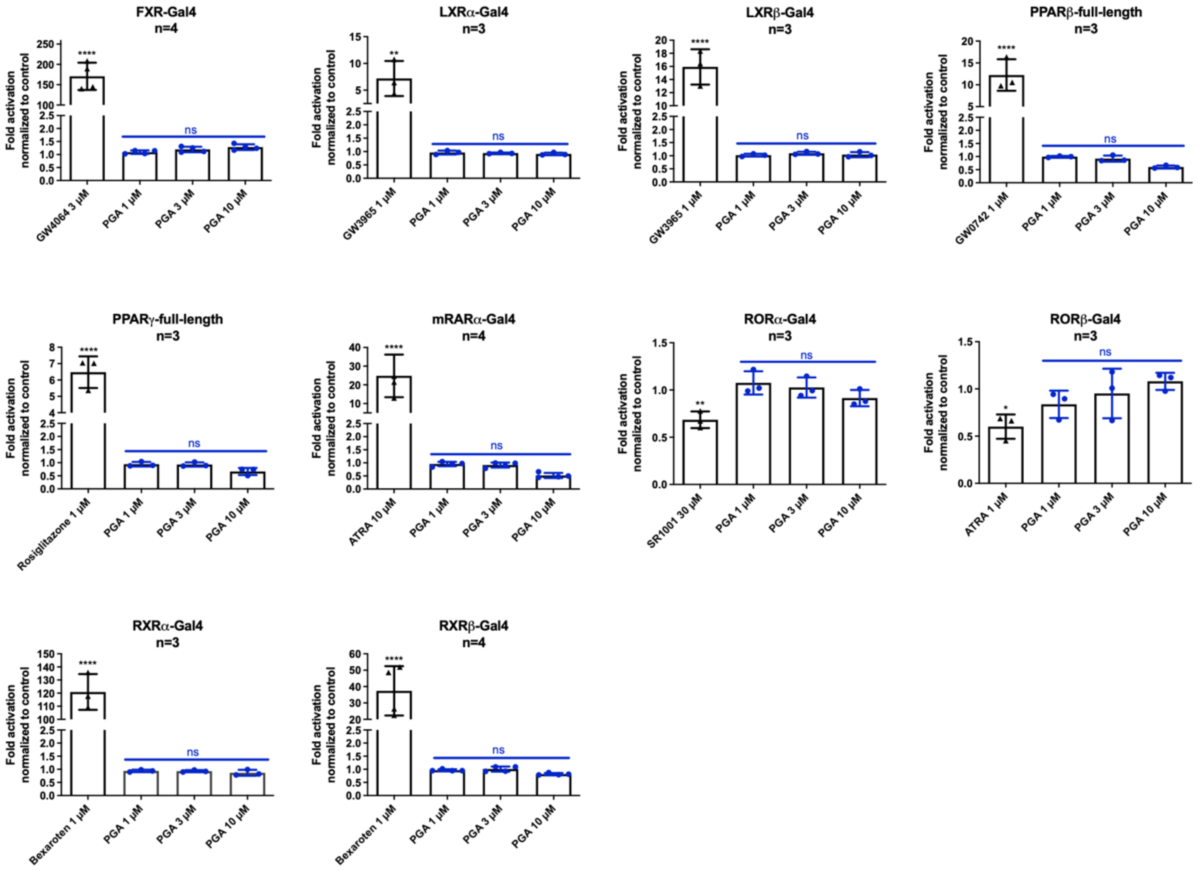


***Figure S 3: PGA does not show off-target effects on a selection of other NRs.*** *NR luciferase assays revealed no off-target effects of PGA on depicted NRs. HEK293 cells were co-transfected with NR-Gal4 (or PPARβ/γ full-length), an UAS luciferase reporter (or a luciferase reporter under the control of PPRE for PPARβ/γ), and eGFP, and treated with PGA at the indicated concentration. The RLU/RFU ratio was measured after an incubation time of 18 hours and normalized to the vehicle control. Mean ± SD, n=4 (FXR-Gal4, mRARα-Gal4, RXRβ-Gal4) or n=3 (rest) in technical quadruplicates. One-way ANOVA with Dunnett's test was performed for statistical analysis. **** P ≤ 0.0001, ** P ≤ 0.01, * P ≤ 0.05, ns P > 0.05.*

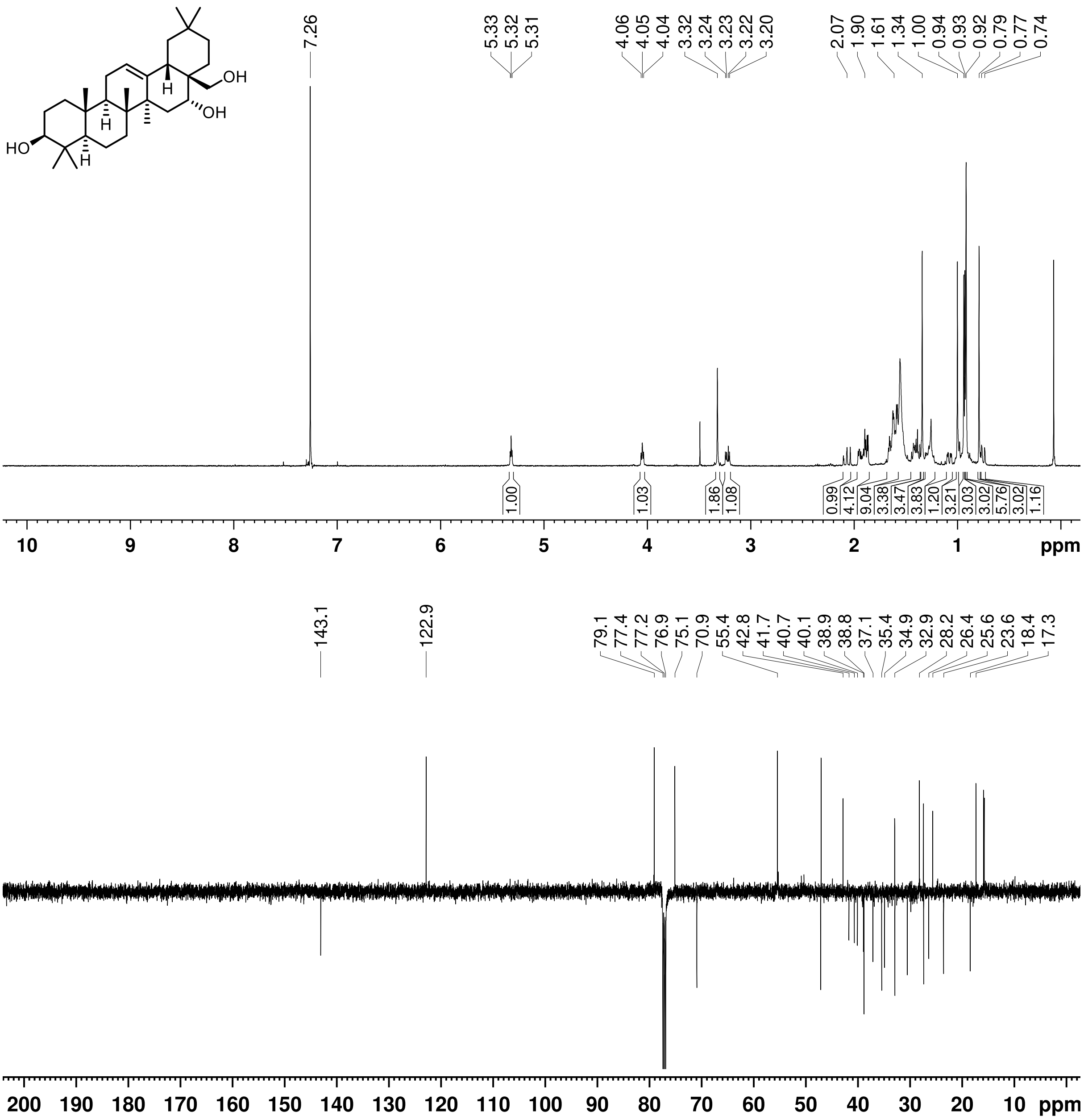


***Figure S 4:*** *[1]H-NMR spectrum (400 MHz, $CDCl_3$; top) and [13]C-NMR (APT) spectrum (126 MHz, $CDCl_3$; bottom) of primulagenin A.*

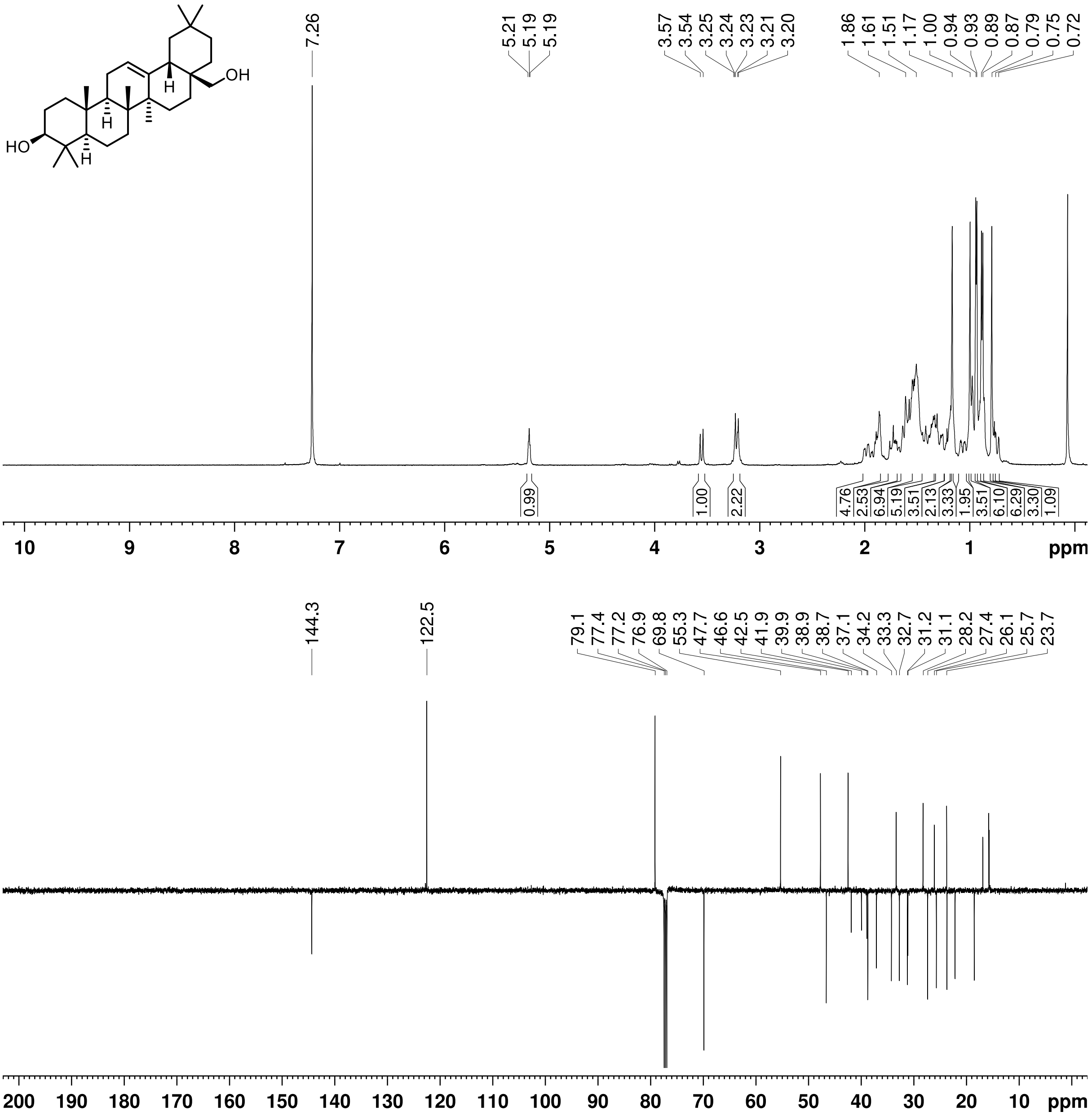


***Figure S 5:*** *[1]H-NMR spectrum (400 MHz, $CDCl_3$; top) and [13]C-NMR (APT) spectrum (126 MHz, $CDCl_3$; bottom) of erythrodiol.*

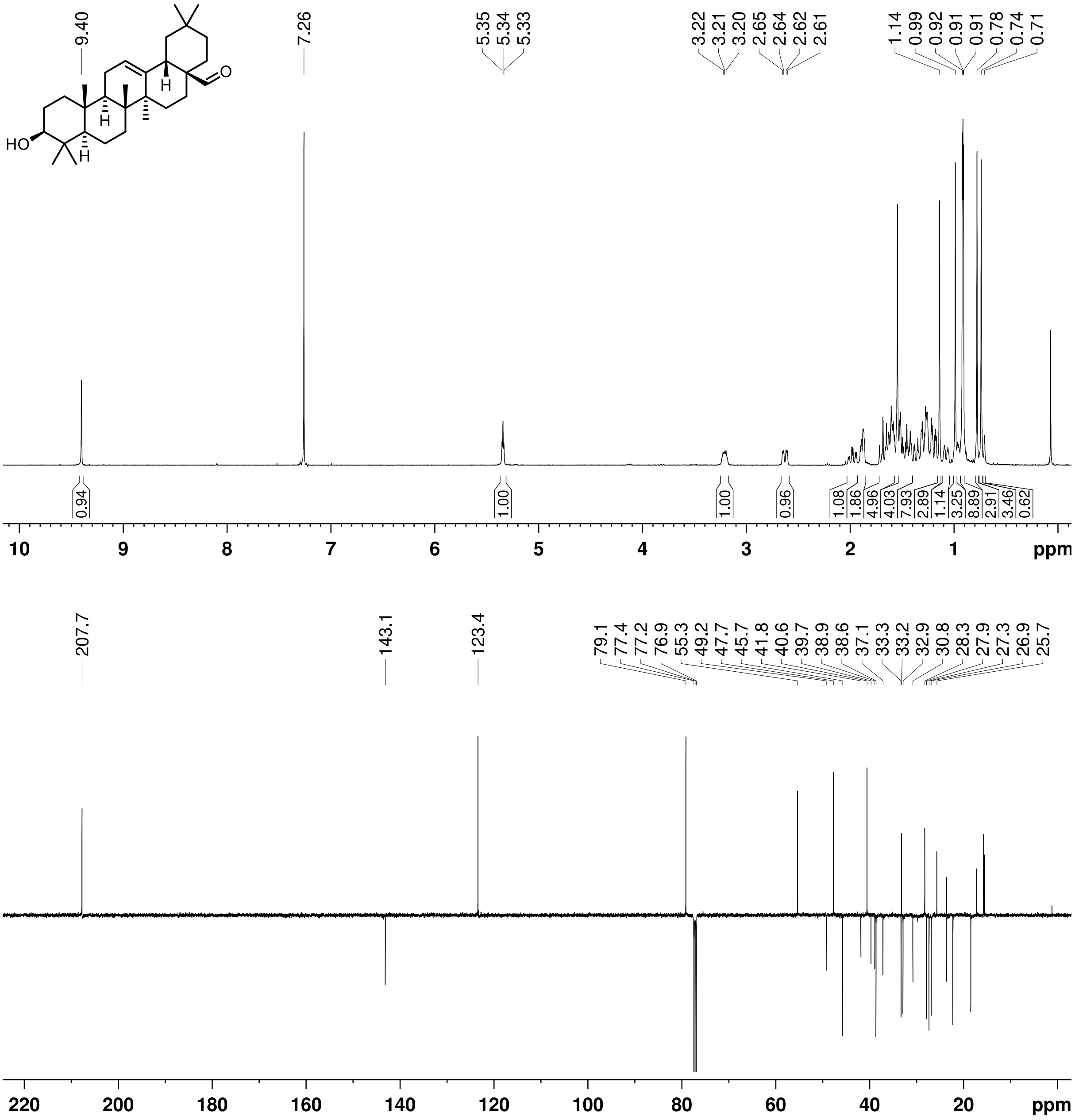


***Figure S 6:*** *$^1$H-NMR spectrum (400 MHz, $CDCl_3$; top) and $^{13}$C-NMR (APT) spectrum (126 MHz, $CDCl_3$; bottom) of oleanolic aldehyde.*

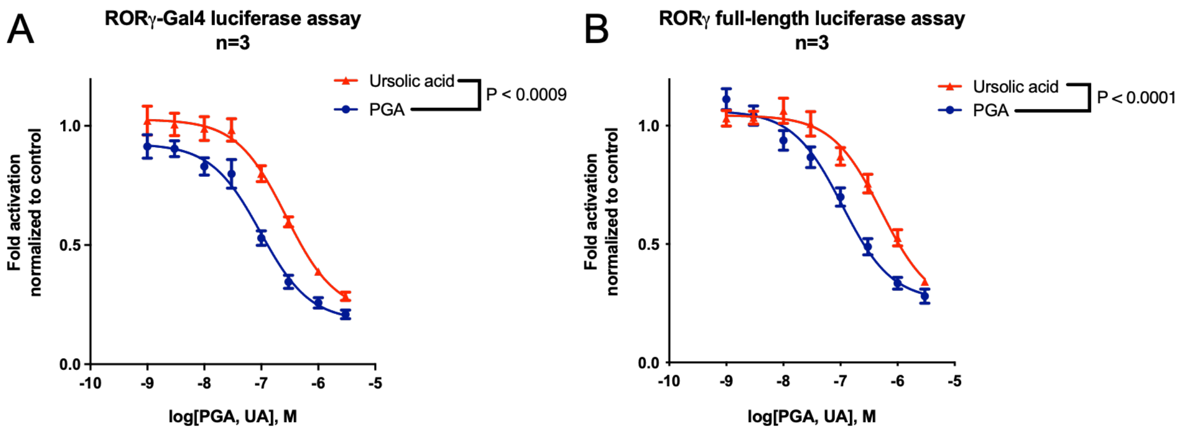


***Figure S 7: PGA shows are higher potency compared to UA. A*** *Concentration-response curves of PGA and UA using a RORγ-Gal4 luciferase assay. HEK293 cells were co-transfected with RORγ-Gal4, an UAS luciferase reporter, and eGFP, and treated with the triterpenoids at the indicated concentration. The RLU/RFU ratio was measured after an incubation time of 18 hours and normalized to the vehicle control. Mean ± SEM, n=3 in technical quadruplicates. An extra sum-of-squares F test was used to statistically compare the best-fit log $IC_{50}$ values of both compounds. P value is depicted in the figure.* ***B*** *Concentration-response curves of PGA and UA using a full-length RORγ luciferase assay. HEK293 cells were co-transfected with full-length RORγ, a luciferase reporter under the control of RORE, and eGFP, and treated with the triterpenoids at the indicated concentration. The RLU/RFU ratio was measured after an incubation time of 18 hours and normalized to the vehicle control. Mean ± SEM, n=3 in technical quadruplicates. An extra sum-of-squares F test was used to statistically compare the best-fit log $IC_{50}$ values of both compounds. P value is depicted in the figure.*

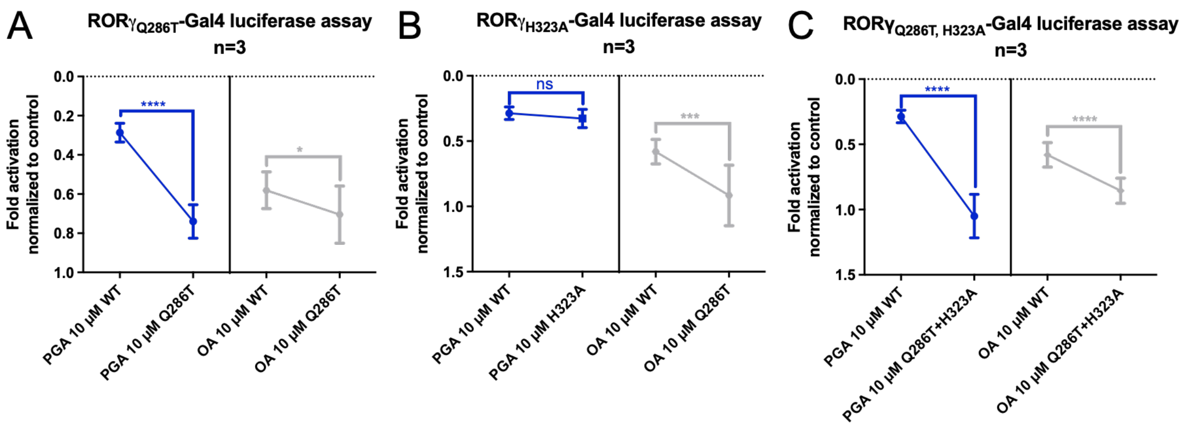


***Figure S 8: PGA and OA are differentially affected by mutations within the RORγ LBD. A-C*** *$RORγ_{mutant}$-Gal4 luciferase assays showing the differences in activity between PGA or OA on the WT NR vs. the respective mutant NR. HEK293 cells were co-transfected with the RORγ-Gal4 WT or a mutant, an UAS luciferase reporter, and eGFP, and treated with PGA or OA at 10 µM. The RLU/RFU ratio was measured after an incubation time of 18 hours and normalized to the vehicle control. Mean ± SD, n=3 in technical quadruplicates. Student's two-tailed t test was performed for statistical analysis. **** P ≤ 0.0001, *** P ≤ 0.001 * P ≤ 0.05, ns P > 0.05.*

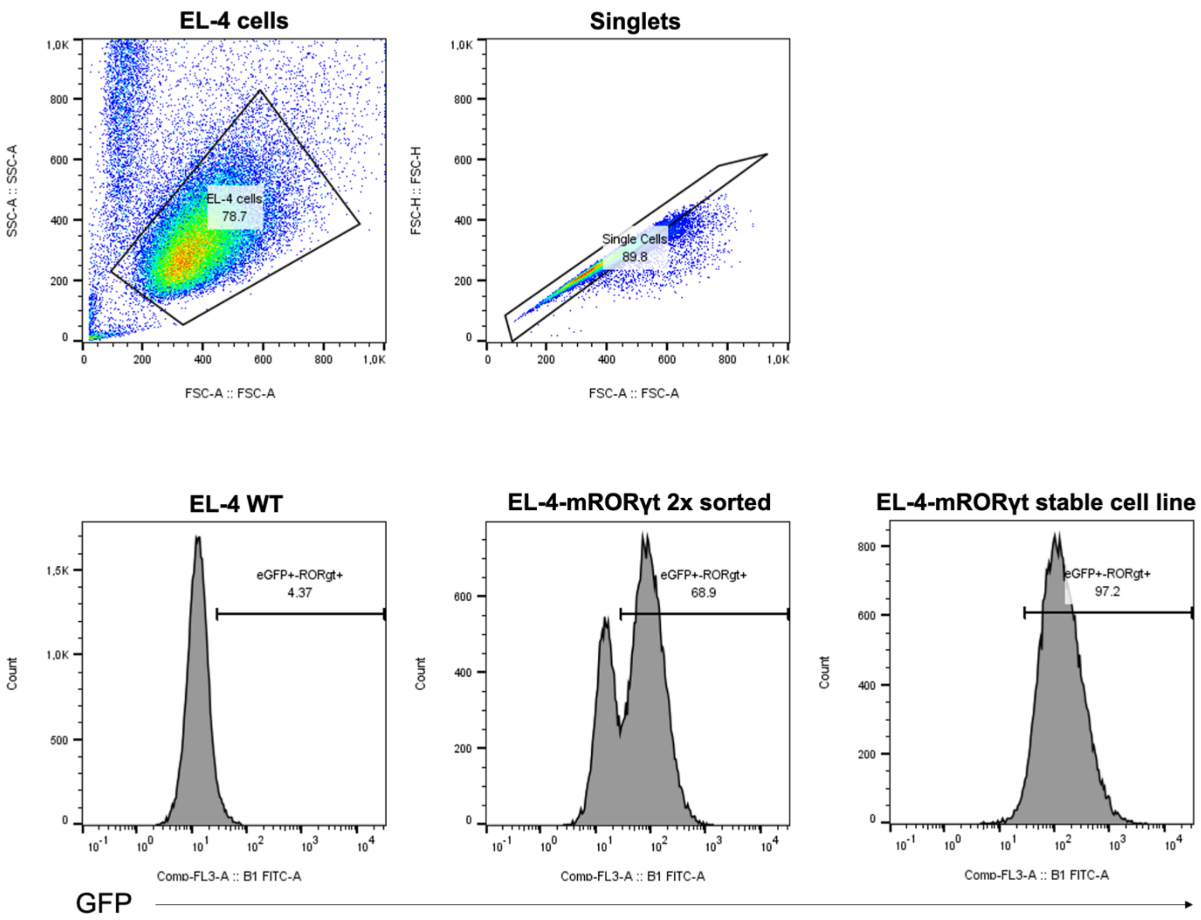


***Figure S 9:*** *Flow cytometry gating strategy for determining the percentage of eGFP⁺ (RORγt⁺) EL-4 cells.*

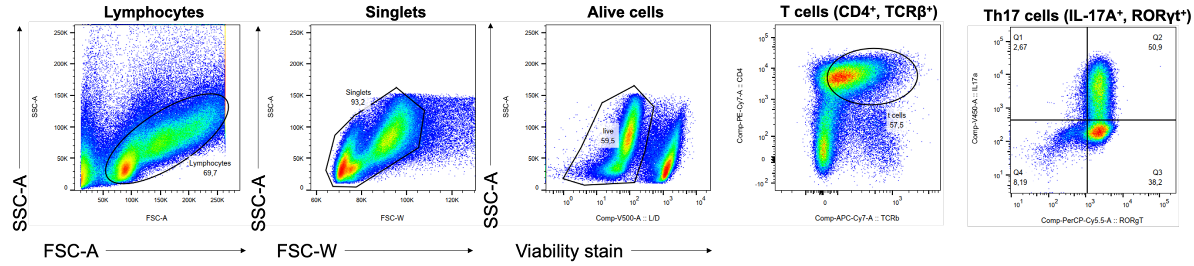


***Figure S 10:*** *Flow cytometry gating strategy for experiments involving murine Th17 cells.*

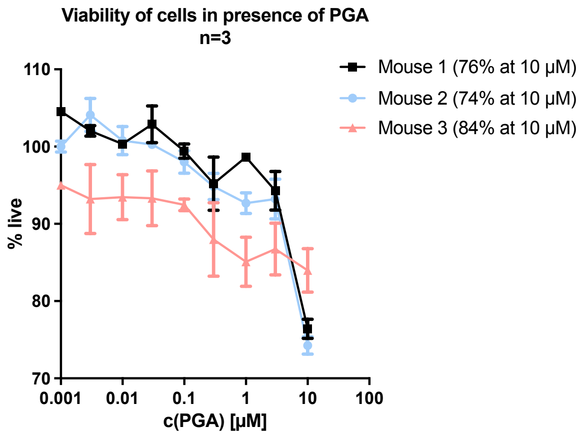


***Figure S 11:*** *Viability of murine OVA-specific CD4⁺ T cells upon PGA treatment. Dead cells were identified using the LIVE/DEAD Fixable Aqua Dead Cell Stain kit (for gating strategy, see* ***Figure S 10****).*

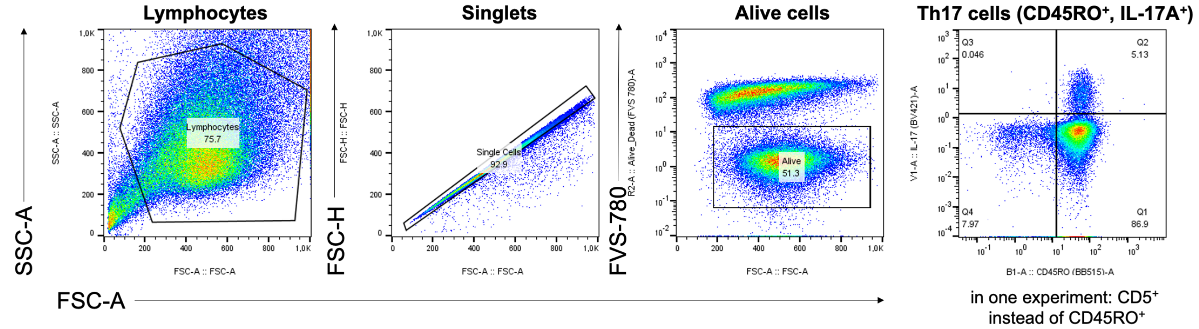


***Figure S 12:*** *Flow cytometry gating strategy for experiments involving human Th17 cells. In one experiment, α-CD5-FITC was used instead of α-CD45RO-BB515.*

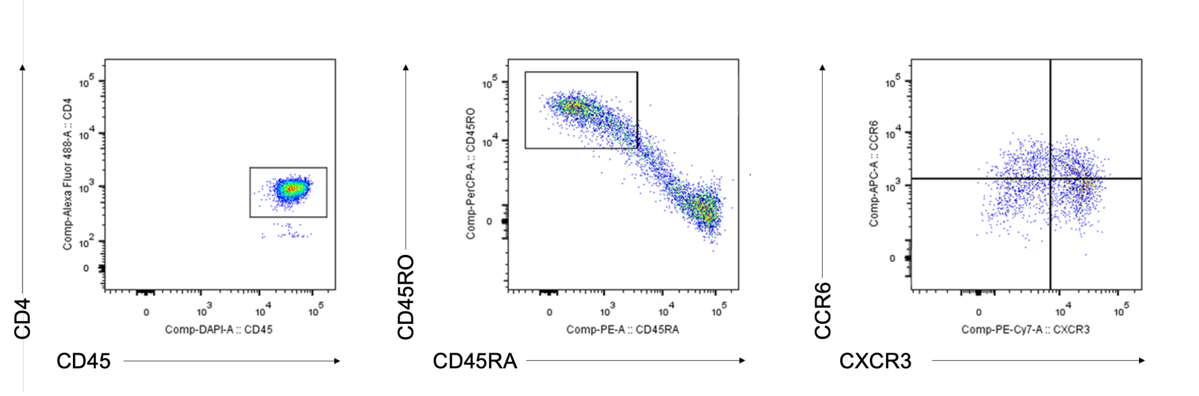


***Figure S 13:*** *FACS strategy for collecting human Th17 cells ($CD45^+CD4^+CD45RA^-CD45RO^+CXCR3^-CCR6^+$ cells).*

***Table S 1:*** *Purity data of triterpenoids that were not commercially obtained. [1]This sample was obtained from the compound library of the Department of Pharmaceutical Sciences, Division of Pharmacognosy (University of Vienna).*

| Triterpenoid | Purity |
|---|---|
| PGA (from the compound library[1])<br>PGA (isolated from Primulae radix)<br>PGA (synthesized) | 91% (UPLC-MS)<br>95% (UPLC-MS) / 97.4% ($^{1}$H-NMR)<br>95.6% (SFC-ELSD) |
| PGD | 95% (UPLC-MS) |
| OAL | 95% (UPLC-MS) |

***Table S 2:*** *Statistical comparison of the best-fit log $IC_{50}$ values of the triterpenoids depicted in* ***Figure 3*** *using an extra sum-of-squares F test. n=3 in all cases.*

| PGA versus… | Null hypothesis | Alternative hypothesis | P value | Conclusion |
|---|---|---|---|---|
| **RORγ-Gal4 luciferase assay** | | | | |
| **PGD** | Log $IC_{50}$ is the same for all data sets. | Log $IC_{50}$ is different for at least one data set. | <0.0001 | Reject null hypothesis. |
| **EA** | | | <0.0001 | |
| **OA** | | | <0.0001 | |
| **β-Amy** | | | <0.0001 | |
| **RORγ full-length luciferase assay** | | | | |
| **PGD** | Log $IC_{50}$ is the same for all data sets. | Log $IC_{50}$ is different for at least one data set. | <0.0001 | Reject null hypothesis. |
| **EA** | | | <0.0001 | |
| **OA** | | | <0.0001 | |
| **β-Amy** | | | 0.0002 | |

***Table S 3:*** *PCR settings for site-directed mutagenesis.*

| Cycle step | Temperature | Time | Cycles |
|---|---|---|---|
| Initial denaturation | 95 °C | 2 minutes | 1 |
| Denaturation | 95 °C | 15 seconds | 18 |
| Primer annealing | 60 °C | 10 seconds | |
| Extension | 68 °C | 3 minutes (30 seconds/bp) | |
| Final synthesis | 68 °C | 5 minutes | 1 |
| Cooling | 4 °C | | 1 |

***Table S 4:*** *PCR settings for amplification of mRORγt and addition of NheI and XhoI restriction sites.*

| Cycle step | Temperature | Time | Cycles |
|---|---|---|---|
| Initial denaturation | 95 °C | 4 minutes | 1 |
| Denaturation | 95 °C | 30 seconds | 29 |
| Primer annealing | 60 °C | 30 seconds | |
| Extension | 72 °C | 1 minute (30 seconds/bp) | |
| Final synthesis | 72 °C | 3 minutes | 1 |
| Cooling | 4 °C | | 1 |

***Table S 5:*** *qPCR settings using the GoTaq Green Master Mix.*

| Cycle step | Temperature | Time | Cycles |
|---|---|---|---|
| Initial denaturation | 95 °C | 2 minutes | 1 |
| Denaturation | 95 °C | 15 seconds | 50-60 |
| Primer annealing / extension | 60 °C | 1 minute (+ plate read) | |
| Melting curve | 55-95 °C | | 1 |
| Cooling | 40 °C | | 1 |

***Table S 6:*** *qPCR settings using the Luna Universal qPCR Master Mix.*

| Cycle step | Temperature | Time | Cycles |
|---|---|---|---|
| Initial denaturation | 95 °C | 1 minute | 1 |
| Denaturation | 95 °C | 15 seconds | 45-55 |
| Primer annealing / extension | 60 °C | 30 seconds (+ plate read) | |
| Melting curve | 55-95 °C | | 1 |
| Cooling | 40 °C | | 1 |

***Table S 7:*** *Purchased items, source, and identifier.*

| Purchased item | Source | Identifier |
|---|---|---|
| β-amyrin | MCE | HY-N2922 (purity > 98%) |
| β-ME | Sigma-Aldrich | 21985023 |
| Agarose | BioCat | AGA500-BCAT |
| Ampicillin | Carl Roth | HP62.1 |
| anti-IFNγ | BioXcell | BE0055 |
| anti-IL-4 | BioXcell | BE0045 |
| AseI | NEB | R0526S |
| ATP | Carl Roth | HN35.2 |
| ATRA | Sigma-Aldrich | R2625 |
| BD Cytofix Fixation Buffer | BD | 554655 |
| BD GolgiPlug Protein Transport Inhibitor | BD | 555029 |
| BD GolgiStop Protein Transport Inhibitor | BD | 554724 |
| BD Perm/Wash Buffer | BD | 554723 |
| BD Pharm Lyse Lysing Buffer | BD | 555899 |
| BD Pharmingen Transcription Factor Buffer Set | BD | 562574 |
| Bexarotene | Sigma-Aldrich | SML0282 |
| BSA | Carl Roth | 8076.3 |
| Cell Activation Cocktail (without Brefeldin A) | BioLegend | 423301 |
| CoA | Sigma-Aldrich | C3019 |
| D-luciferin | SynChem | BC218 |
| Digitonin | Sigma-Aldrich | D141 |
| DMEM with phenol red | Sigma-Aldrich | D6546 |
| DMEM without phenol red | Lonza | 12-917F |
| DMSO | Carl Roth | 4720.2 |
| DTT | Sigma-Aldrich | 43815 |
| EasySep Human CD4+ T Cell Isolation Kit | Stemcell Technologies | 100-0696 |
| Echinocystic acid | MCE | HY-N0271 (purity > 98%) |
| Erythrodiol | MCE | HY-N2419 (purity > 98%) |
| eGFP-N1 | Clontech | 6085-1 |
| EL-4 cells | ECACC | 85023105 |
| EMEM | Lonza | 12-125F |
| FBS | Biowest | S1810; batch number S00CN |
| FCS (for mTh17 cells) | Gibco | A5256801 |
| G418 | Sigma-Aldrich | G8168-10ML |
| *G6PC* primer | Qiagen | 249900 |
| *GAPDH* primer | Qiagen | 249900 |
| *GlutaMAX* | Gibco | 35050038 |
| GoTaq Green Master Mix | Promega | M712 |
| GW0742 | Sigma-Aldrich | G3295 |
| GW3965 | Sigma-Aldrich | G6295 |
| GW4064 | Sigma-Aldrich | G5172 |
| HEK293 cells | ATCC | CRL-1573 |

| | | |
|---|---|---|
| HepG2 cells | ATCC | HB-8065 |
| Herculase II Fusion DNA Polymerase | Agilent | 600677 |
| High-Capacity cDNA Reverse Transcription Kit | Thermo Fisher Scientific | 4368814 |
| IL-1β (for mTh17 cells) | PeproTech | 211-11B |
| IL-1β (human) | Miltenyi Biotec | 130-093-897 |
| IL-6 | PeproTech | 216-16B |
| IL-23 (for mTh17 cells) | R&D | 1887-ML |
| IL-23 (human) | Miltenyi Biotec | 130-095-757 |
| innuPREP RNA Mini Kit 2.0 | IST Innuscreen | 845-KS-2040010 |
| Ionomycin | Sigma-Aldrich | I9657 |
| IPTG | Euromedex | EU0008-C |
| $KH_2PO_4$ | Carl Roth | 3904.1 |
| L-glutamine | Lonza | BE17-605E |
| LB | Carl Roth | X964.2 |
| Lipofectamine 3000 | Thermo Fisher Scientific | L3000001 |
| Lipofectamine LTX Reagent with PLUS Reagent | Thermo Fisher Scientific | 12343593 |
| Luna Universal qPCR Master Mix | NEB | M3003 |
| Lymphopure | BioLegend | 426202 |
| MeOH | VWR | 83638.320 |
| Monarch DNA Gel Extraction Kit | NEB | T1020S |
| Monensin | BioLegend | 420701 |
| $Na_2$- EDTA*$2H_2O$ | Carl Roth | 8043.2 |
| $Na_2HPO_4$ | Carl Roth | P030.2 |
| NaCl | Carl Roth | 3957.2 |
| Naive $CD4^+$ T Cell Isolation Kit II, human | Miltenyi Biotec | 130-094-131 |
| NheI-HF | NEB | R3131S |
| Oleanolic acid | Sigma-Aldrich | O5504 (purity ≥97%) |
| OVA peptide | AnaSpec | AS27025 |
| Penicillin–Streptomycin | Lonza | DE17-602E |
| Penicillin-Streptomycin (for mTh17 cells) | Gibco | 15140122 |
| pIRES2-eGFP | Clontech | 6029-1 |
| PMA | Sigma-Aldrich | P1585 |
| Primulae radix | Kottas | Batch number: P21301312; voucher specimen JR-20210422_A1 deposited in the Herbarium of the Department of Pharmaceutical Sciences, Division of Pharmacognosy |
| PureLink HiPure Plasmid Midiprep Kit | Thermo Fisher Scientific | K210005 |
| QuikChange Lightning Site-Directed Mutagenesis Kit | Agilent | 210519 |
| Reporter Lysis 5X Buffer | Promega | E4030 |
| Resazurin | Sigma-Aldrich | 199303 |
| Rosiglitazone | Sigma-Aldrich | R2408 |
| RPMI 1640 | Lonza | 12-167F |
| RPMI 1640 (for mTh17 cells) | Gibco | 31870025 |
| SR1001 | Sigma-Aldrich | SML0322 |

| | | |
|---|---|---|
| SR2211 | Sigma-Aldrich | SML1170 |
| SYBR Safe DNA Gel Stain | Thermo Fisher Scientific | S33102 |
| T0901317 | Tocris Bioscience | 2373 |
| T4 DNA Ligase | NEB | M0202S |
| TCEP | Hampton Research | HR2-651 |
| TGF-β | R&D | 240-B-002 |
| Thrombin | MP Biomedicals | 02154163-CF |
| Trypsin | Thermo Fisher Scientific | 27250-018 |
| Ursolic acid | Sigma-Aldrich | 89797 (purity ≥98.5%) |
| Whole blood | Austrian Red Cross | Donor #1: A0040220902051<br>Donor #2: A004022092582U<br>Donor #3: A004022119347L |
| XhoI | NEB | R0146S |
| XL1-Blue Competent Cells | Agilent | 200249 |
| Zeocin | Thermo Fisher Scientific | J67140.8EQ |

***Table S 8:*** *Antibodies and viability dye used for experiments involving murine Th17 cells.*

| Antibody | Fluorophore | Species | Isotype | Clone | Company | Identifier |
|---|---|---|---|---|---|---|
| CD4 | Pe-Cy7 | Mouse | IgG2b, kappa | GK1.5 | Thermo Fisher Scientific | 25-0041-82 |
| TCR-β | APC-Cy7 | Mouse | IgG | H57-597 | Thermo Fisher Scientific | 47-5961-82 |
| IL-17A | V450 | Mouse, Rat | IgG2a, kappa | eBio17B7 | Thermo Fisher Scientific | 48-7177-82 |
| RORγt | PCP-Cy5.5 | Mouse | IgG2a, kappa | Q31-378 | BD | 562663 |
| Viability dye: LIVE/DEAD Fixable Aqua Dead Cell Stain Kit | | | | | Thermo Fisher Scientific | L34957 |

***Table S 9:*** *Antibodies and viability dye used for experiments involving human Th17 cells.*

| Antibody | Fluorophore | Species | Isotype | Clone | Company | Identifier |
|---|---|---|---|---|---|---|
| CCR6 | Alexa Fluor 647 | Human | Mouse IgG2b, kappa | G034E3 | BioLegend | 353403 |
| CD3 | / | Human | Mouse (BALB/c) IgG1, kappa | UCHT1 | BD | 555329 |
| CD4 | FITC | Human | Mouse / IgG2b, kappa | OKT-4 | Thermo Fisher Scientific / Invitrogen | 11-0048-42 |
| CD5 | FITC | Human | Mouse IgG1, kappa | UCHT2 | BD | 555352 |
| CD45 | Brilliant Violet 421 | Human | Mouse IgG1, kappa | 2D1 | BioLegend | 368521 |
| CD45RA | PE | Human | Mouse IgG2b, kappa | HI100 | BioLegend | 304107 |
| CD45RO | BB515 | Human | Mouse (BALB/c) IgG2a, kappa | UCHL1 | BD | 564529 |
| CD45RO | PerCP/Cyanine5.5 | Human | Mouse IgG2a, kappa | UCHL1 | BioLegend | 304221 |
| CXCR3 | PE/Cyanine7 | Human | Mouse IgG1, kappa | G025H7 | BioLegend | 353719 |
| IL-17A | BV-421 | Human | Mouse IgG1, kappa | N49-653 | BD | 562933 |
| Viability dye: BD Horizon Fixable Viability Stain 780 | | | | | BD | 565388 |

***Table S 10:*** *List of gifted plasmids and their providers used in this work. [1]The Herbert Wertheim UF Scripps Institute for Biomedical Innovation & Technology, University of Florida; [2]Department of Pharmacy, LMU Munich; [3]Department of Biochemistry, Nihon University School of Medicine; [4]Center for Integrative Genomics, University of Lausanne; [5]Institute of Genetics, Molecular and Cellular Biology; [6]Salk Institute for Biological Studies; [7]Department of Laboratory Medicine, Medical University of Vienna*

| Plasmid | Provider |
|---|---|
| hRORα-Gal4 | Prof. Laura A. Solt[1] |
| hRORβ-Gal4 | |
| hRORγ-Gal4 | |
| hRORγ (full-length) | Prof. Patrick Griffin[1] |
| mRORγt (full-length) | Prof. Laura A. Solt[1] |
| FXR-Gal4 | Prof. Daniel Merk[2] |
| LXRα-Gal4 | Prof. Makoto Makishima[3] |
| LXRβ-Gal4 | |
| PPARβ (full-length) | Prof. Walter Wahli[4] |
| PPARγ (full-length) | |
| mRARα-Gal4 | Prof. Hinrich Gronemeyer[5] |
| RXRα-Gal4 | Prof. Ronald Evans[6] |
| RXRβ-Gal4 | Prof. Hinrich Gronemeyer[5] |
| UAS-LUC | Prof. Ronald Evans[6] |
| RORE-LUC | Prof. Patrick Griffin[1] |
| PPRE-LUC | Prof. Nikolina Papac[7] |

***Table S 11:*** *List of primers used in this work.*

| Primer name | Forward primer (5'-3') | Reverse primer (5'-3') | Comments |
|---|---|---|---|
| **Quik RORgGal4 Gln286Thr** | ACA GGG AGA CAT GCA CGC TGC GGC TGG AGG | CCT CCA GCC GCA GCG TGC ATG TCT CCC TGT | Mutagenic primer used to generate the hRORγ-Gal4 Q286T mutant and the Q286T+H323A double mutant. |
| **hRORg-Gal4 H323A** | ACG GTG TGC CCA CGC CCT CAC CGA GGC C | GGC CTC GGT GAG GGC GTG GGC ACA CCG T | Mutagenic primer used to generate the hRORγ-Gal4 H323A mutant. |
| **GAL4-BD** | TCA TCG GAA GAG AGT AG | / | Standard sequencing primer from Microsynth used to verify the Q286T, H323A, and Q286+H323A mutations. |
| **mRORgt pIRES2-eGFP NheI fwd / XhoI rev** | AAA GCT AGC GGT GGA ATA CCA TGA GAA CAC | AAA CTC GAG TCG AGT CAC TTT GAC AGC CCC | Primers for PCR-amplification of mRORγt and addition of NheI and XhoI restriction sites. |
| **IRES-for** | TAG GCG TGT ACG GTG GG | / | Standard sequencing primer from Microsynth used to verify the mRORγt-IRES-GFP construct. |
| ***mil17a*** | CAA CCG TTC CAC GTC ACC C | GAG CTT CCC AGA TCA CAG AGG G | qPCR primer. |
| ***mil17f*** | GAA ACC AGC ATG AAG TGC ACC C | TGC TAC CTC CCT CAG AAT GGC | qPCR primer. |
| ***mil23r*** | AGG CTT TTC GGA ACC TCA TGC | GTC AGA TTG CTG GGG GCA TC | qPCR primer. |
| ***m18s*** | GTA ACC CGT TGA ACC CCA TT | CCA TCC AAT CGG TAG TAG CG | qPCR primer. |
| ***IL-17A*** | ACC GAT CCA CCT CAC CTT GG | AGT CCA CGT TCC CAT CAG CG | qPCR primer. |
| ***G6PC*** | Hs_G6PC_1_SG QuantiTect Primer Assay (GeneGlobe Id: QT00031913; detected transcript: NM_000151) | | qPCR primer. |
| ***GAPDH*** | Hs_GAPDH_1_SG QuantiTect Primer Assay (GeneGlobe Id: QT00079247; detected transcript: NM_001256799) | | qPCR primer. |
